# Beta-caryophyllene enhances wound healing through multiple routes

**DOI:** 10.1101/611046

**Authors:** Sachiko Koyama, Anna Purk, Manpreet Kaur, Helena A. Soini, Milos V. Novotny, Keith Davis, C. Cheng Kao, Hiroaki Matsunami, Anthony Mescher

**Affiliations:** Department of Biology, Biotechnology Program, Indiana University, Bloomington, IN, USA; School of Public Health, Indiana University, Bloomington, IN, USA; Department of Psychological and Brain Sciences, Indiana University, Bloomington, Indiana, USA; Department of Chemistry, and Institute for Pheromone Research, Indiana University, Bloomington, Indiana, USA; Department of Molecular and Cellular Biochemistry, Indiana University, Bloomington, Indiana, USA; Department of Molecular Genetics and Microbiology, School of Medicine, Duke University, Durham, North Carolina, USA; Department of Anatomy and Cell Biology, School of Medicine, Indiana University, Bloomington, Indiana, USA

## Abstract

Beta-caryophyllene is an odoriferous bicyclic sesquiterpene found in various herbs and spices. Recently, it was found that beta-caryophyllene is a ligand of the cannabinoid receptor 2 (CB2). Activation of CB2 will decrease pain, a major signal for inflammatory responses. We hypothesized that beta-caryophyllene can affect wound healing by decreasing inflammation. Here we show that cutaneous wounds of mice treated with beta-caryophyllene had enhanced re-epithelialization. The treated tissue showed increased cell proliferation and cells treated with beta-caryophyllene showed enhanced cell migration, suggesting that the higher re-epithelialization is due to enhanced cell proliferation and cell migration. The treated tissues also had up-regulated gene expression for hair follicle bulge stem cells. Olfactory receptors were not involved in the enhanced wound healing. Transient Receptor Potential channel genes were up-regulated in the injured skin exposed to beta-caryophyllene. Interestingly, there were sex differences in the impact of beta-caryophyllene as only the injured skin of female mice had enhanced re-epithelialization after exposure to beta-caryophyllene. Our study suggests that chemical compounds included in essential oils have the capability to improve wound healing, an effect generated by synergetic impacts of multiple pathways.

## Introduction

Odors from other conspecific^1^individuals [1] and the environment [2-6] affect the behaviors and physiological status in many animals. The significance of olfactory stimuli tends to be underestimated in case of humans. However, we have a long history of actively utilizing odorants to heal or alter our physiological conditions. Various herbal plants’ and spices’ extracts have been used to reduce stress, pain, and to promote recovery from injury or illness [2-6]. In spite of such a long history of using herbal plant extracts and although an increasing number of scientists are starting to test natural products owing to their diverse molecular scaffolds and biologically active substructures, there is still a strong need to examine the influences of extracts. It is especially important to better understand their mechanisms of action.

When essential oils are used instead of single chemical compound, there have been challenges to examine their mechanism of action because of the large variances in the percentage each chemical compound present [7] depending on the extraction methods [8], the area of origin including differences in the geographic altitude of the areas [9], seasons the plants were harvested [10], and parts of the plants extracted [11, 12]. It is thus important to utilize chemically-pure compounds or chemically-defined extracts as described in some recent studies [13-17]. These studies enable testing the roles of individual chemicals and obtaining clear mechanistic insights [13, 14].

In addition, the route of administration can affect the efficacy of the odorants. Odorants of herbal plants are often called “aromas”, which suggests that the route is through the olfactory system. However, there are various routes that odorant chemicals can exert effects. First, olfactory receptors are expressed in both olfactory neurons and in non-olfactory tissues, such as skin and circulatory organs [18, 19]. Furthermore, some odorants activate both olfactory receptors and non-olfactory receptors. For example, beta-caryophyllene (BCP), which is an odoriferous bicyclic sesquiterpene present in many herbs and spices, is a ligand of cannabinoid receptor 2 (CB2)^2^ [20] as well as olfactory receptors. How odorants such as BCP act on distinct receptor classes is important for the most effective use of BCP.

The CB2 receptor is expressed in neuronal cells, immune tissue, hair follicles, sebaceous glands, the dermomuscular layer, and vascular smooth muscle in intact skin [21]. Activation of CB2 by CB2 selective agonist GP1a improved re-epithelialization in wound healing [22, 23]. Essential oil of *Copaifera paupera,* which contains BCP also improved wound healing [24], although it is not clear whether BCP had a direct or indirect role and whether the wound healing is promoted by the CB2 receptor. In this project, we examined whether BCP can improve re-epithelialization and, if so, whether the olfactory system is involved in the impact.

## Results and discussion

### BCP treatment enhanced re-epithelialization

Wound healing is a series of overlapping cellular and biochemical processes that restore the integrity of injured tissues. In general, healing occurs with four overlapping phases that takes place from minutes to weeks: hemostasis, inflammation, cell proliferation and migration, and maturation/scarring [25-28]. We examined whether exposure to BCP affect cell proliferation and the re-epithelialization of mouse skin in a full thickness wound generated on the back of mice using scissors. The wound was then covered with a device that can retain either 50 uL of BCP (50mg/kgbw, diluted with olive oil) or a control (olive oil alone) (see Materials and Methods for details, S1 Fig). Throughout the study, none of the mice exhibited behaviors suggestive of irritation with the BCP (S2, S3 Fig).

Immunofluorescence staining of keratin-14 (K14), a marker of re-epithelialization [29], revealed that wounds treated with BCP had increased keratinocyte migration distance from intact skin near the wound toward the wound center relative to the controls, indicating improved re-epithelialization (Fig. 1). Staining for proliferating cell nuclear antigen (PCNA) showed that many cells were proliferating at the wound edge in both the BCP and Oil groups (green color in the wound edge area in Fig. 1b). In the BCP group, PCNA+ fibroblast cells are concentrated in the reticular dermis (lower area of dermis). The fibroblast cells in the reticular dermis migrate into the wound bed and form an extracellular matrix of granulation tissue [30]. The results on K14 and PCNA show that the BCP-treated group exhibits higher re-epithelialization and cell proliferation.

**Fig 1.**
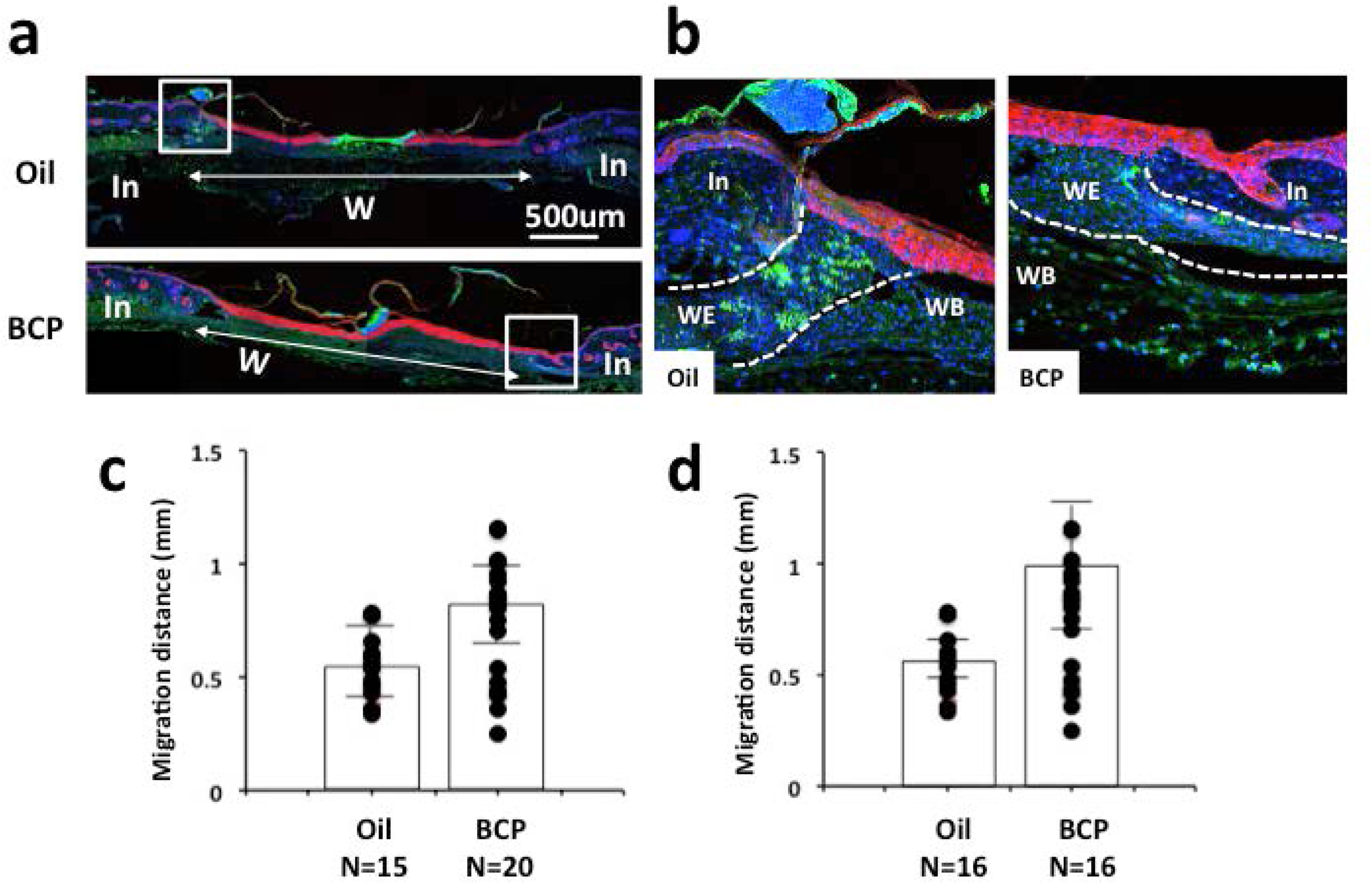
Re-epithelialization measured by keratin 14. **(a)** Immunofluorescence staining of K14 (red) and PCNA (green) with nuclear staining using Draq5 (blue) of skin with wound treated with BCP or with oil. **(b)** Higher magnification images of areas of boxes in **(a)**. PCNA was localized in the nucleus and, when density of cells increased, it became localized in the cytoplasm **(b)**. This was observed at the center of the wound. In: intact area, W: wounded area. **(c, d)** Length of K14+ staining area from the boundary of intact and wounded area to the center of the wound in skin harvested on post-surgery day 3 **(c)** and 4 **(d)**. Dots indicate each datum and the height of bars indicates the median. Horizontal lines indicate quartiles. 3^rd^day, Kolmogorov-Smirnov test, *P*=0.008, BCP, n=20, Oil, n=15; 4^th^day, Kolmogorov-Smirnov test, *P*=0.002, *: *P*<0.05, BCP, n=16, Oil, n=16.

Another marker of re-epithelialization is the expression of filaggrin, a marker of differentiated skin [31]. Filaggrin is expressed in the upper layer of the epidermis (stratum corneum) and functions as a skin barrier, protecting the body from losing water and from exogenous pathogens [31]. In the BCP-treated wounds, the range of differentiated stratum corneum layer in the invading epidermis was wider and the expansion from the wound edge was longer when compared to the Oil group (Fig. 2 red).

**Fig. 2.**
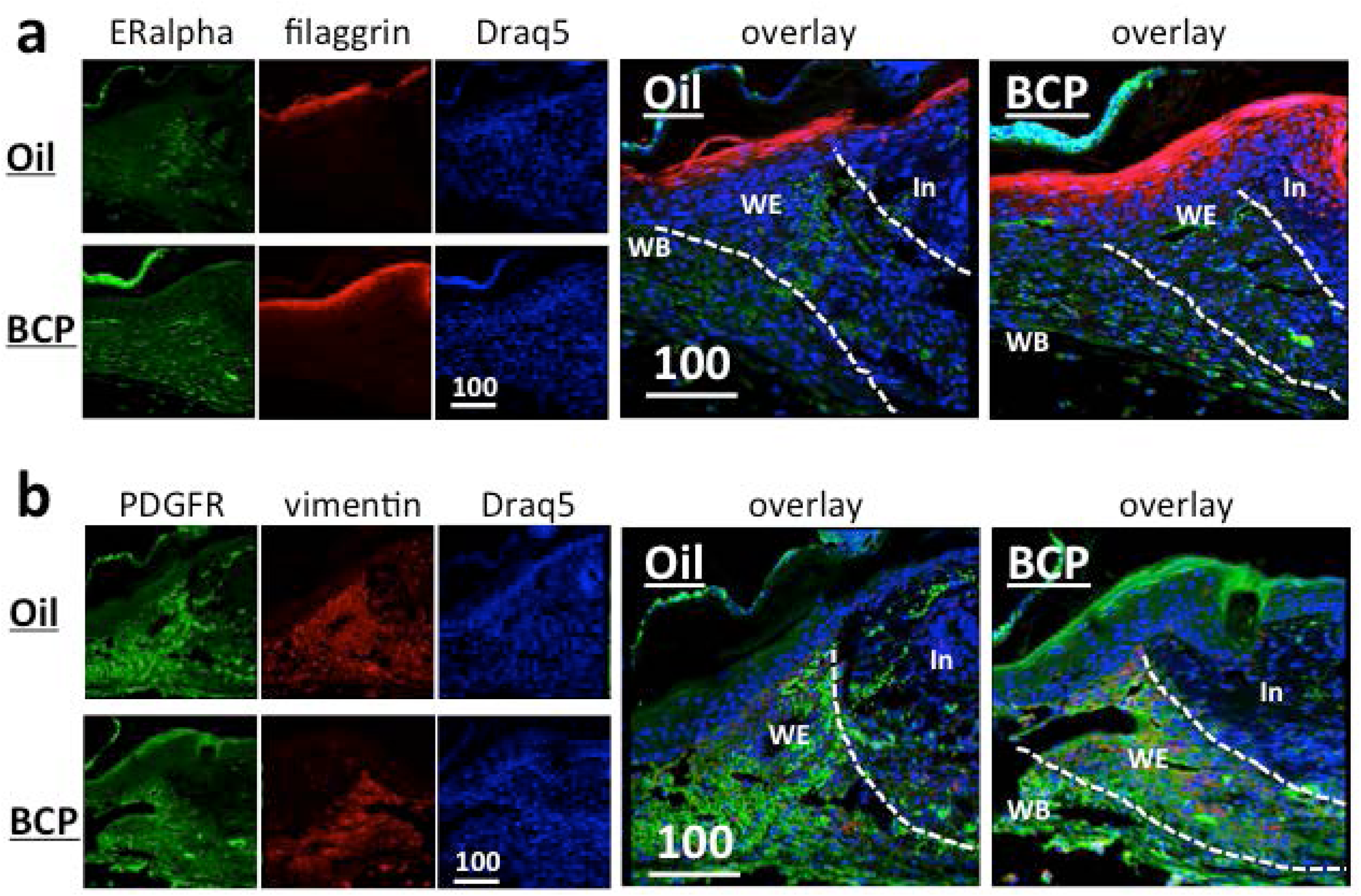
Expression of filaggrin (red) and estrogen alpha (green) (a), and PDGFR (green) and vimentin (red) (b) in the wounded area of mice. **(a)** View at the wound edge. In: intact area, WE: wound edge, WB: wound bed. Scale bar is in μm. Dotted lines show the boundaries between the intact area and wound edge, and the wound edge and wound bed. Filaggrin staining of the BCP group was broader in width and longer in the distance from the wound edge. Estrogen alpha staining did not show obvious differences between groups. **(b)** PDGFR expression in the epidermis was stronger in the BCP group. Both the BCP and Oil group showed a large area of staining of PDGFR in the wound edge (WE) and wound bed (WB). Vementin staining did not show obvious differences between the BCP group and Oil group. In: intact area, WE: wound edge, WB: wound bed. Scale bars are in μm.

Estrogen is known to accelerate wound healing [32]. We examined if there are differences in the expression of estrogen receptor alpha after exposure to BCP. Estrogen receptor alpha was expressed strongly in the dermis fibroblasts of both the BCP and Oil group without differences (Fig. 2).

Platelet-derived growth factor-A and –B (PDGF-A and –B) are expressed in fibroblasts and keratinocytes [33]. PDGF receptors (PDGFR) are expressed in fibroblasts, smooth muscle cells, epithelial cells, macrophages, and their expression increases when there is inflammation [33]. In wound healing, PDGF stimulates mitogenesis and chemotaxis of fibroblasts, smooth muscle cells, neutrophils and macrophages, and stimulates production of fibronectin, collagen, proteoglycans and hyaluronic acid [33]. PDGFRα also has a role in both extracellular matrix and hepatocyte growth factor production in fibroblasts in remodeling of connective tissue [34]. Immunofluorescence staining for PDGFRα showed that it is expressed strongly in the epidermis and the wound edge of the tissue treated with BCP (Fig. 2b). In the tissues treated with Oil, PDGFRα was expressed strongly only at the wound edge (Fig. 2b). Vimentin, which has a role in fibroblast proliferation and keratinocyte differentiation [35, 36], was expressed strongly in the wound edge (Fig. 2b red) of both the BCP-treated and Oil-treated wound. These results suggest that the impact of BCP is primarily on the invasion of epidermis and differentiation of keratinocytes, resulting in enhance re-epithelialization of the wounded area.

### BCP treatment enhanced cell proliferation

To determine the rate and location of cell proliferation in the wounded tissue, 5-bromo-2’-deoxyuridine (BrdU) was injected twice every 2 hours and tissues were harvested 2 hours after the second injection (see Materials andMethods for details). Significantly more BrdU+ cells were found on the basal layer of the interfollicular epidermis and hair follicles, and in the papillary dermis [37-39] in the BCP-treated wounds. In both the BCP-treated and controls, there were many BrdU+ cells at the wound margin and wound bed (Fig. 3). Previous studies have shown that proliferation takes place at a proliferative hub, which is from 2 mm outside of the wound edge until 3 mm from the wound edge [40]. We observed that cell proliferation in the controls took place highly at the wound margin and at the basal layer of the interfollicular epidermis (Fig. 3f, Oil). In contrast, the BCP-treated wounds had higher cell proliferation in the interfollicular epidermis, in the dermis, at the hair follicles, and in the wound bed (Fig. 3f, BCP).

**Fig 3.**
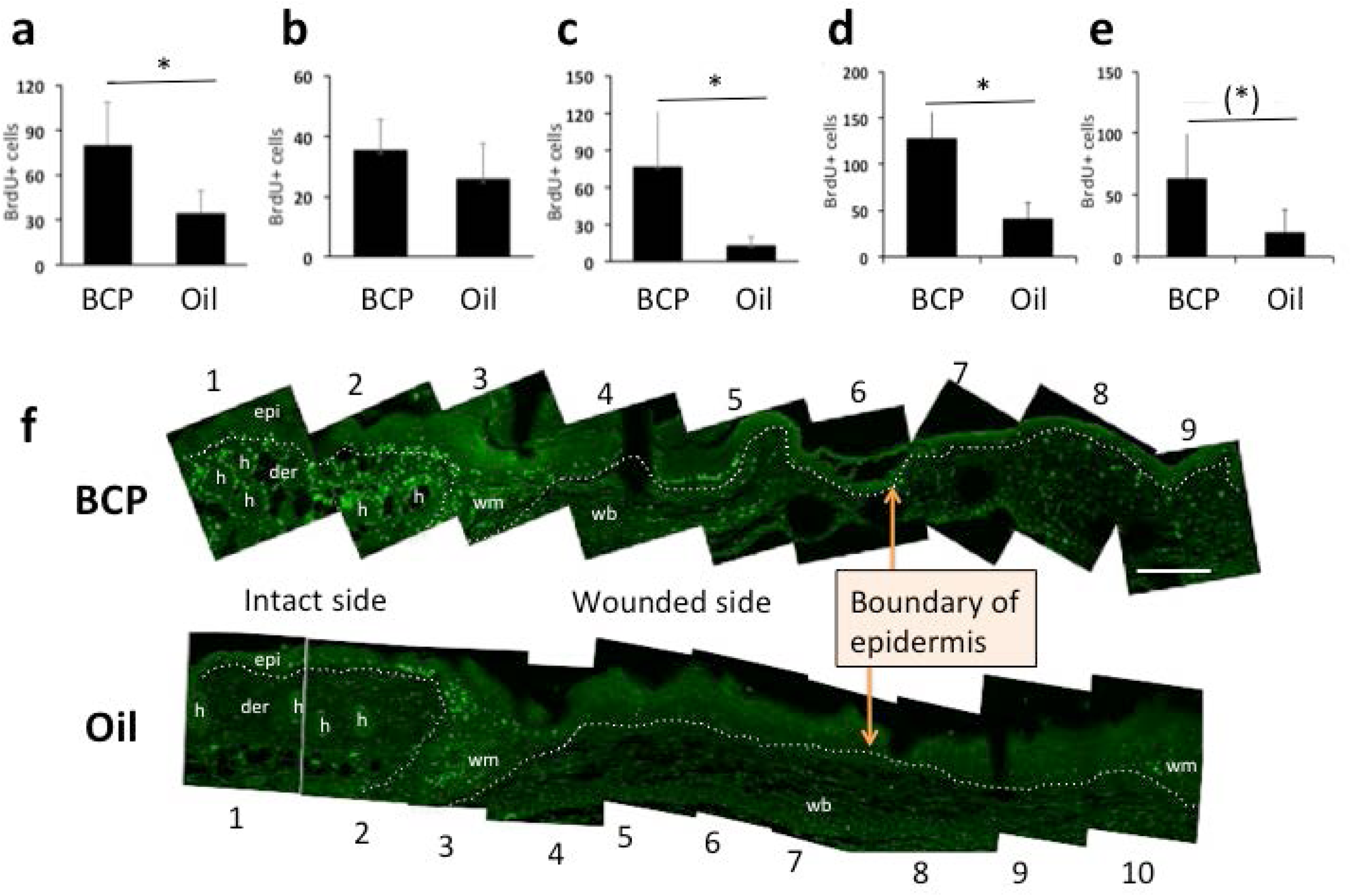
BrdU+ cells in the epidermis and dermis. **(a)** BrdU+ cells in the epidermis (epi in **(f)**) (Student T-test, T=3.077, df=8, *P*=0.015, BCP, n=5, Oil, n=5), **(b)** BrdU+ cells in the wound margin (wm in **(f)**) (Student T-test, T=1.343, df=8, *P*=0.216, BCP, n=5, Oil, n=5), **(C)** BrdU+ cells at the hair follicle (h in **(f)**) (Student T-test, T= 3.038, df=8, *P*=0.015, BCP, n=5, Oil, n=5). **(d)** BrdU+ cells at the dermis excluding wound margin (Student T-test, T=4.645, df=8, *P*=0.002, BCP, n=5, Oil, n=5). **(e)** BrdU+ cells at the BrdU+ cells at the wound bed (Student T-test, T=2.197, df=8, *P*=0.059, BCP, n=5, Oil, n=5). **(f)** Representative views of combined photos used in quantification for each group. h: hair follicle, epi: epidermis, der: dermis, wm: wound margin, wb: wound bed, scale bar: 200 µm. Numbers below and above the photos are operationally assigned numbers for explanation in the text on locations in the wound. Dotted lines show the boundaries between epidermis and dermis, dermis and wound margin zone, and wound margin zone and wound bed.

Stem cells in the hair follicle contribute to re-epithelialization by migrating from the hair follicle bulge to epidermis and then to the center of the wound [41]. The strong BrdU staining of hair follicles in the BCP-treated wounds compared to the controls suggest that a larger number of hair follicle stem cells may have contributed to the enhanced re-epithelialization in BCP group.

If enhanced re-epithelialization was due to a reduction of inflammation and an early shift to the cell proliferation/migration stage, then BCP should not impact cultured cells. To test this hypothesis, we conducted time-lapse imaging for 7 hours and enumerated cell division. We conducted this with different concentrations of BCP. We found a peak of cell proliferation at 26 µM BCP, which decreased at higher concentrations (Fig. 4). In higher concentrations we observed less cell proliferation. These results suggested that BCP treatment was on cell proliferation and enhanced re-epithelialization rather than the suppression of inflammation.

**Fig 4.**
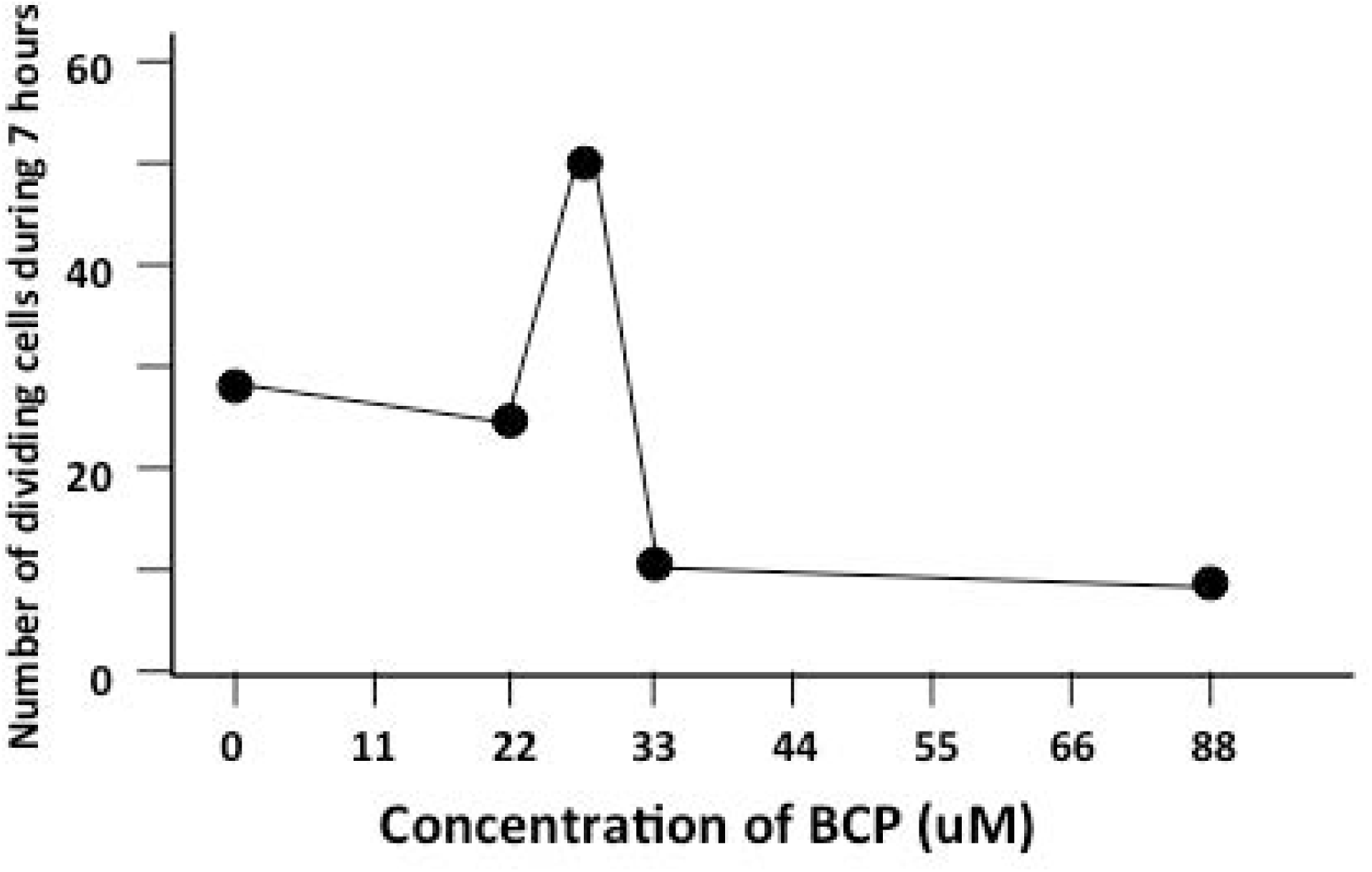
Proliferating primary fibroblast cells exposed to BCP at various concentrations. Number of dividing and apoptotic cells during 7 hours of exposure to BCP.

### BCP enhanced cell migration

The increased cell migration caused by BCP was not reported previously and we examined this further. To determine whether exposure to BCP can stimulate cell migration, we conducted *in vitro* assays of scratch tests and chemotaxis assays. Primary cultured fibroblasts and keratinocytes from C57BL/6 mice exposed to BCP (diluted with DMSO) for 24 hours showed higher chemotactic responses relative to the controls (fibroblasts, 2.1 times higher number of cells, and keratinocytes, 2.5 times higher number of cells, both compared to control condition cells exposed to DMSO) (Fig. 5a, b, c).

**Fig 5.**
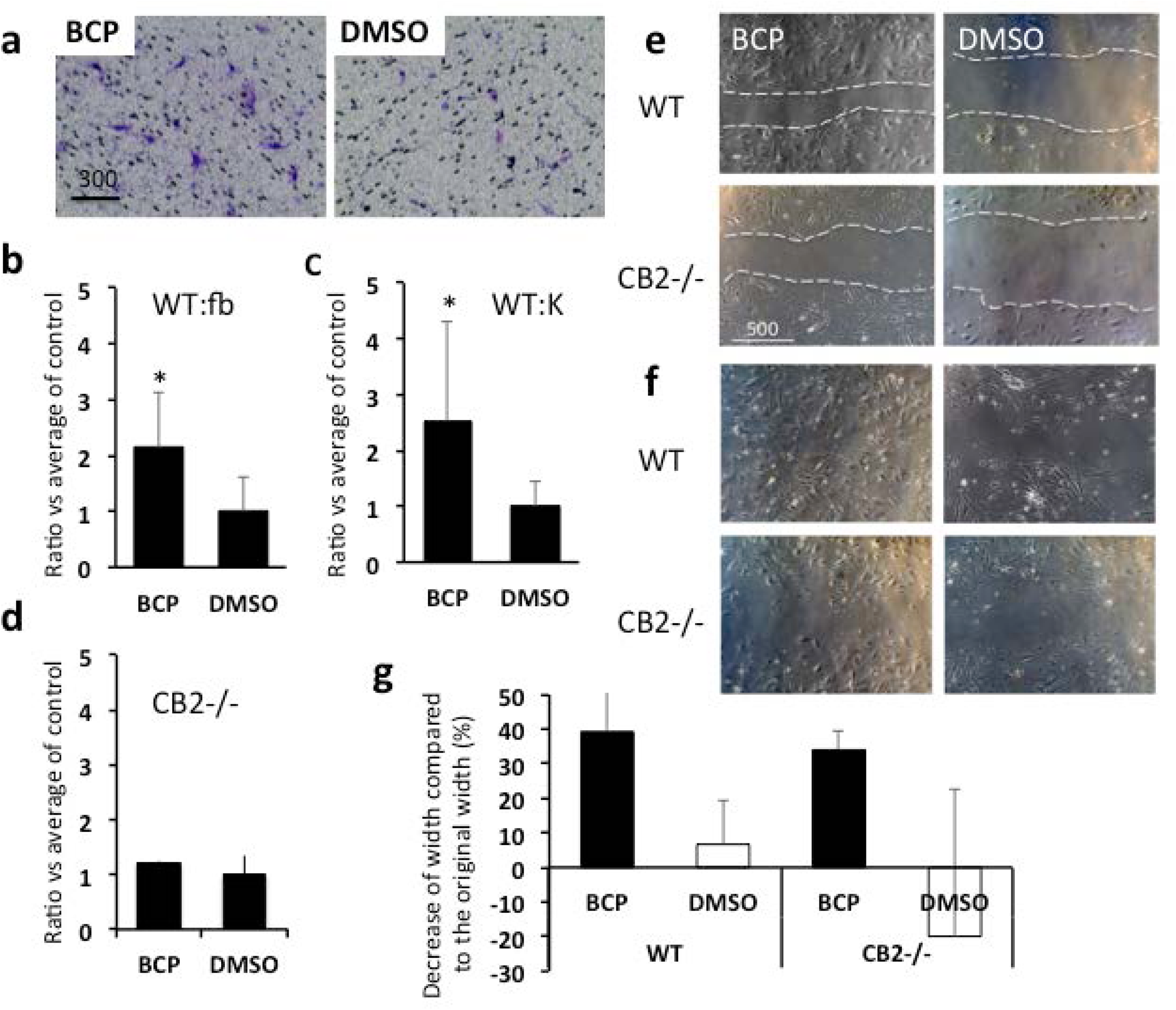
Influence of BCP on cell migration. **(a)** Representative results of chemotaxis assay. Cells showed chemotactic responses to BCP. Fibroblasts and keratinocytes were exposed either to BCP or to DMSO (control) in chemotaxis assay. WT cells exposed to BCP showed significantly more chemotactic responses toward the bottom of the inserts (fibroblasts, Student T-test, T=2.564, df=8, *P*=0.033, BCP, n=4, DMSO, n=4, **(b)**; keratinocytes, Student T-test, T=2.456, df=8, *P*=0.04, BCP, n=4, DMSO, n=4, **(c)**), but cells isolated from CB2 knockout mice did not show differences between those exposed to BCP and those exposed to DMSO control (fibroblasts, Student T-test, T=0.944, df=4, *P*=0.398, BCP, n=3, DMSO, n=3, **(d)**). **(e, f, g)** Results of scratch tests. Representative images on the influence of BCP (27 µM) on cell migration of fibroblasts. Six hours after scratch **(e)** and 1 day after scratch **(f)**. Dotted white lines show the edge of scratched area. Scale bar = µm, **(g)** The distance cells migrated during the 6 hours shown as a comparison to the original width (% of shrink in width). O% means no change from original width. ANOVA, Medium type, *F*_1,8_=7.16, *P*=0.028, cell type, *F*_1,8_=0.365, *P*=0.562, BCP, n=3, DMSO, n=3.

However, exposure to BCP did not stimulate chemotactic responses in fibroblasts isolated from CB2 knockout mice (1.2 times higher number of cells compared to control condition cells exposed to DMSO) (Fig. 5d). These results suggest that activation of CB2 could lead to an increase in chemotactic responses.

In the chemotaxis assay above we found that cells show chemotactic responses toward BCP. We then conducted scratch tests to determine if exposure to BCP stimulates chemotactic responses to repopulate in petri dishes. After culture dish reached confluency, the culture media was replaced to culture media containing BCP (23µM/10 µL DMSO/10 mL cell culture media) or DMSO (10 µL/10mL cell culture media). Exposure to BCP increased cell migration to the scratched area (39.2% decrease in the width in 6 hours), relative to the control (6.7% decrease in the width in 6 hours). The cells from CB2 knockout mice exposed to BCP also showed a comparable rate of repopulating the devoided area (34.0% decrease in the width in 6 hours) (Fig. 5e, f, g). These results suggest that BCP can stimulate cell migration, but not through CB2 pathway.

We started this study with the hypothesis that BCP will positively impact wound healing by activating signaling by CB2. While BCP did improve re-epithelialization, the results of the cell migration studies showed that BCP’s effect on re-epithelialization may be more complicated than through the activation of CB2. Therefore, we examined whether CB2 antagonists and agonists will affect re-epithelialization. AM630, an antagonist of CB2 [42] was daily injected 20 min before the daily application of BCP, CB2 agonist JWH133 [43] was topically applied instead of BCP to determine if it generates a similar impact as BCP. Then tissues of the BCP group, the Oil group, the AMP630+BCP group, and the JWH133 group were harvested on the 5^th^day post-surgery. Sections were stained with K14 antibody and the distance of migration from the edge of the wound was measured. Consistent with our previous observations, topical application of BCP enhanced re-epithelialization. CB2 agonist JWH133 significantly enhanced re-epithelialization compared to the Oil group (Fig. 6). When CB2 antagonist AM630 was injected daily into mice treated with BCP, the results were not clear. Re-epithelialization was not statistically different from the Oil group but the variance was large and there was a trend towards differences (*P*=0.071) (Fig. 6). These results suggest that there could be some other pathways involved in the BCP enhanced re-epithelization of the wounds.

**Fig. 6.**
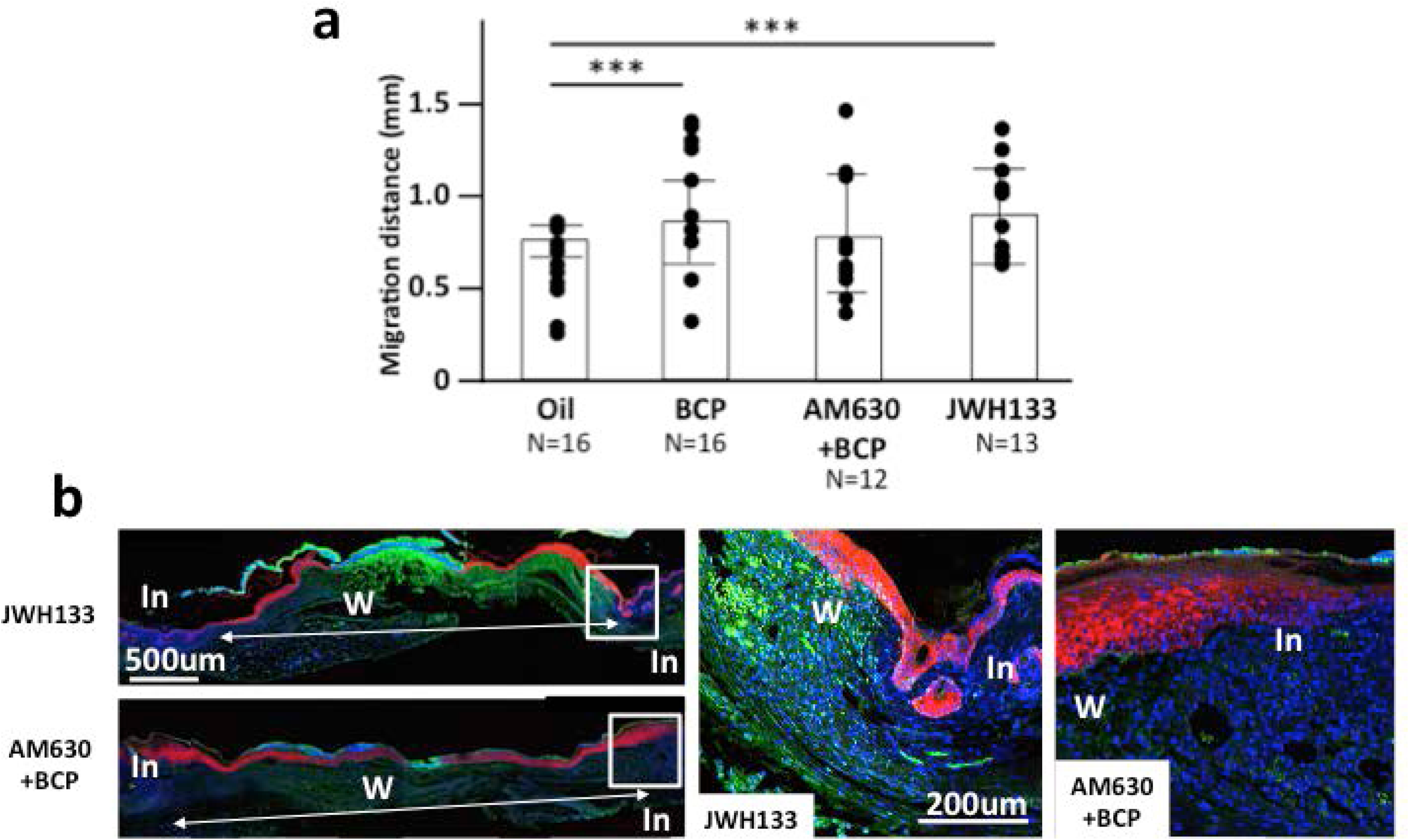
Re-epithelialization (K14+ distance from wound edge) in mice exposed to BCP, Oil, CB2 agonist JWH133, and BCP+CB2 antagonist AM633. **(a)** Length of K14+ staining area from the boundary of the intact and wounded area to the center of the wound in skin harvested on post-surgery day 5 from mice exposed to BCP, oil, JWH133 or injected CB2 antagonist AM630 daily before daily treatment with BCP. Dots indicate each data set and the height of bars indicates the median. Horizontal lines indicate quartiles. ANOVA, *F*_3,53_=5.51, *P*=0.002) (ANOVA, *F*_3,53_=5.51, *P*=0.002. Post-hoc pairwise analyses between BCP vs. oil, Tukey’s post-hoc pairwise comparison, *P*=0.002, CB2 agonist JMW133 group vs. oil group, Tukey’s post-hoc pairwise comparison, *P*=0.071, and CB2 antagonist AM630+BCP group vs. BCP group, Tukey’s pairwise comparison, *P*=0.097; BCP, n=9, Oil, n=10, JWH133, n=8, AM630+BCP, n=7). **(b)** Representative images of JWH133 group and BCP+AM630 group. In: Intact area, W: wounded area.

### BCP changes gene expression in the wounded skin

In order to better predict the pathways through which BCP acts, we conducted RNA sequencing and transcriptome analyses. The skin from mice 17 to 18 hours post-injury treated with BCP or oil was used. The transcriptome of mice that were not injured (NT group) was also analyzed to determine which gene expression changes are caused by injury itself.

When a comparison was made between the BCP group vs. the NT group and the Oil group vs. the NT group, the gene expression patterns of the BCP group and the Oil group were similar (38 out of 50 top significant genes were common). This suggests that these common genes could be genes that become modulated when there is skin injury (S5 Fig).

When comparison was made between the BCP group vs. the Oil group, significant differences in gene expression between the BCP group and the Oil group were found (Fig 7). BCP-treatment significantly up-regulated a large number of genes, the 50 whose expression changed the most are in Table 1. Twenty percent of the up-regulated genes code for keratins (*Krt*) or keratin-associated proteins (*Krtap*) [44] (Table 1). Krtap genes are expressed in cells of the hair shaft cortex [45], where we had observed an increased number of BrdU+ cells (Fig. 3). Other than the possible role of BCP on enhancing re-epithelialization, this result also suggests the possibility that exposure to BCP promotes more complete skin regeneration and hair neogenesis. The remaining forty of the top 50 up-regulated genes code for functions in cell migration (e.g. *Adamts*) [46], cell fate determination, and hair follicle formation (*Bambi, Msx2, Dlx3, Padi1, Hoxc13, S100a*) [47-52]. These results overall suggest that BCP has impacts on hair follicle stem cell production and skin regeneration to aid in wound healing.

**Fig 7.**
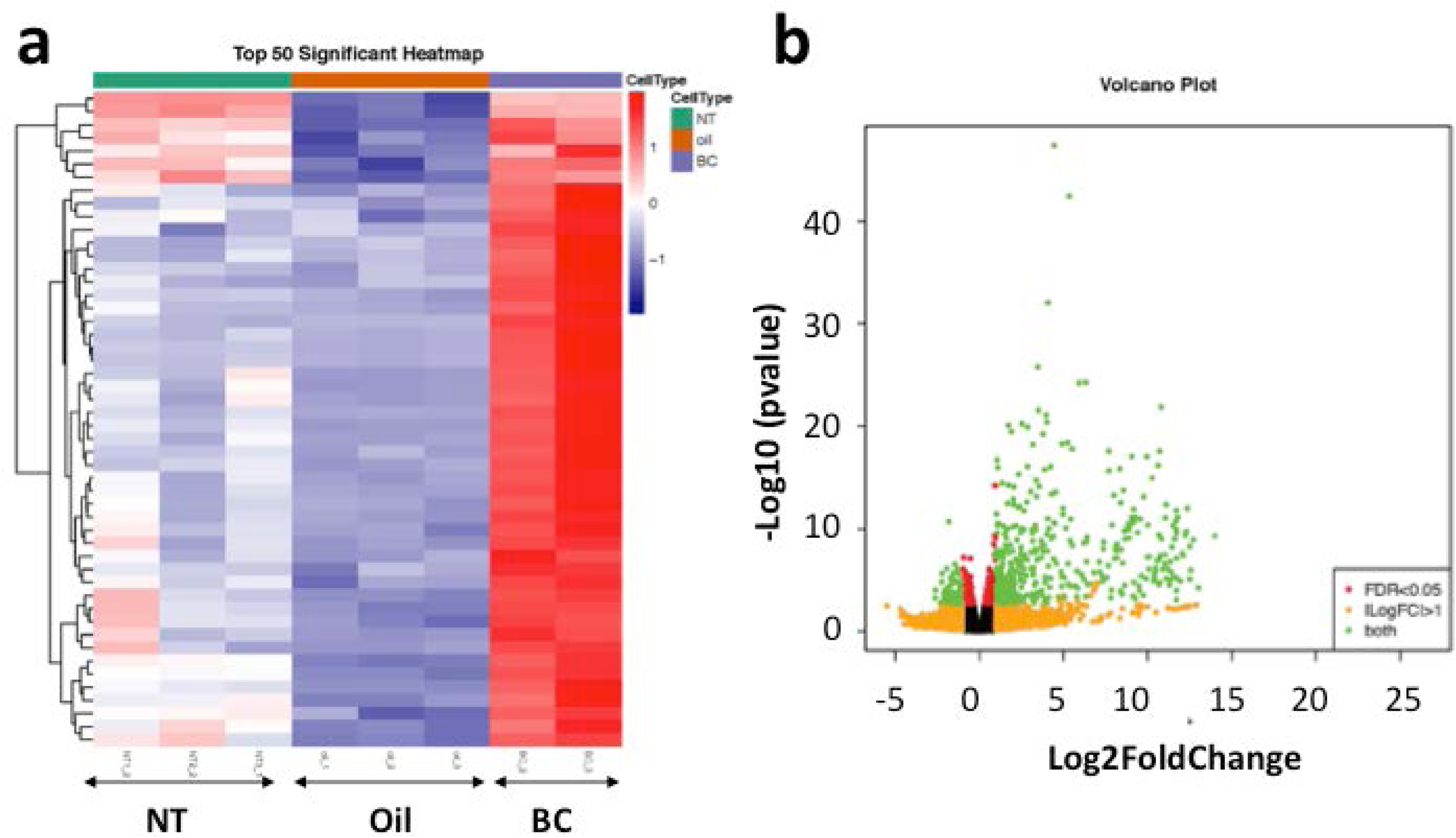
Results of RNA sequencing of post-surgery 17 hours skin and intact skin. Heatmap showing the top 50 significant gene expressions in the skin exposed to BCP (n=2) or oil (n=3), 17 to 18 hours post-surgery (inflammation stage), and in the skin of mice without skin excision (NT group) (n=3). Comparison between BCP vs. oil **(a)** showed a clear difference between BCP and Oil. **(b)** Volcano plot from the comparison between BCP vs. Oil. Y-axis shows log10 (p value) and X-axis shows log2 fold change value.

**Table 1.**
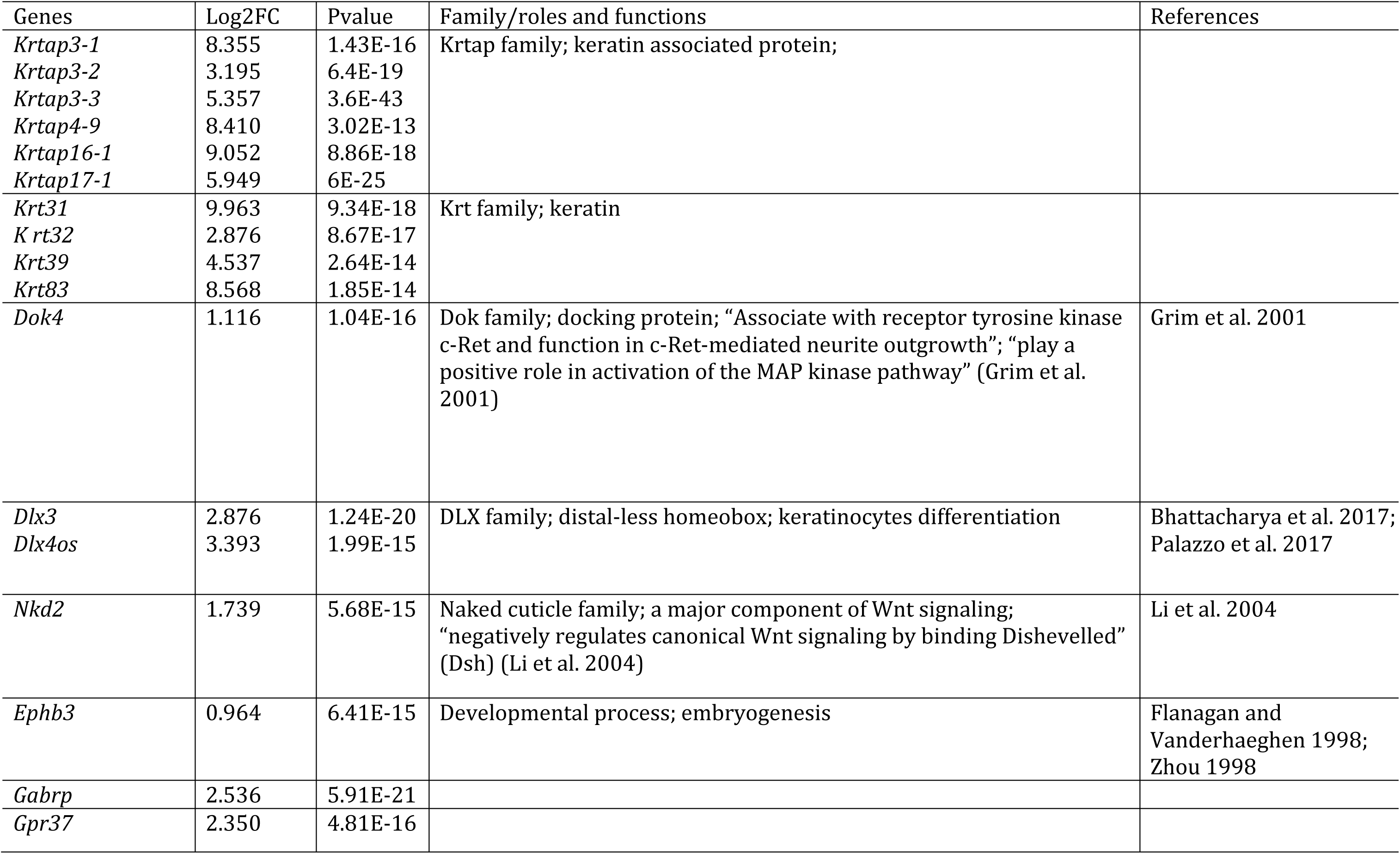
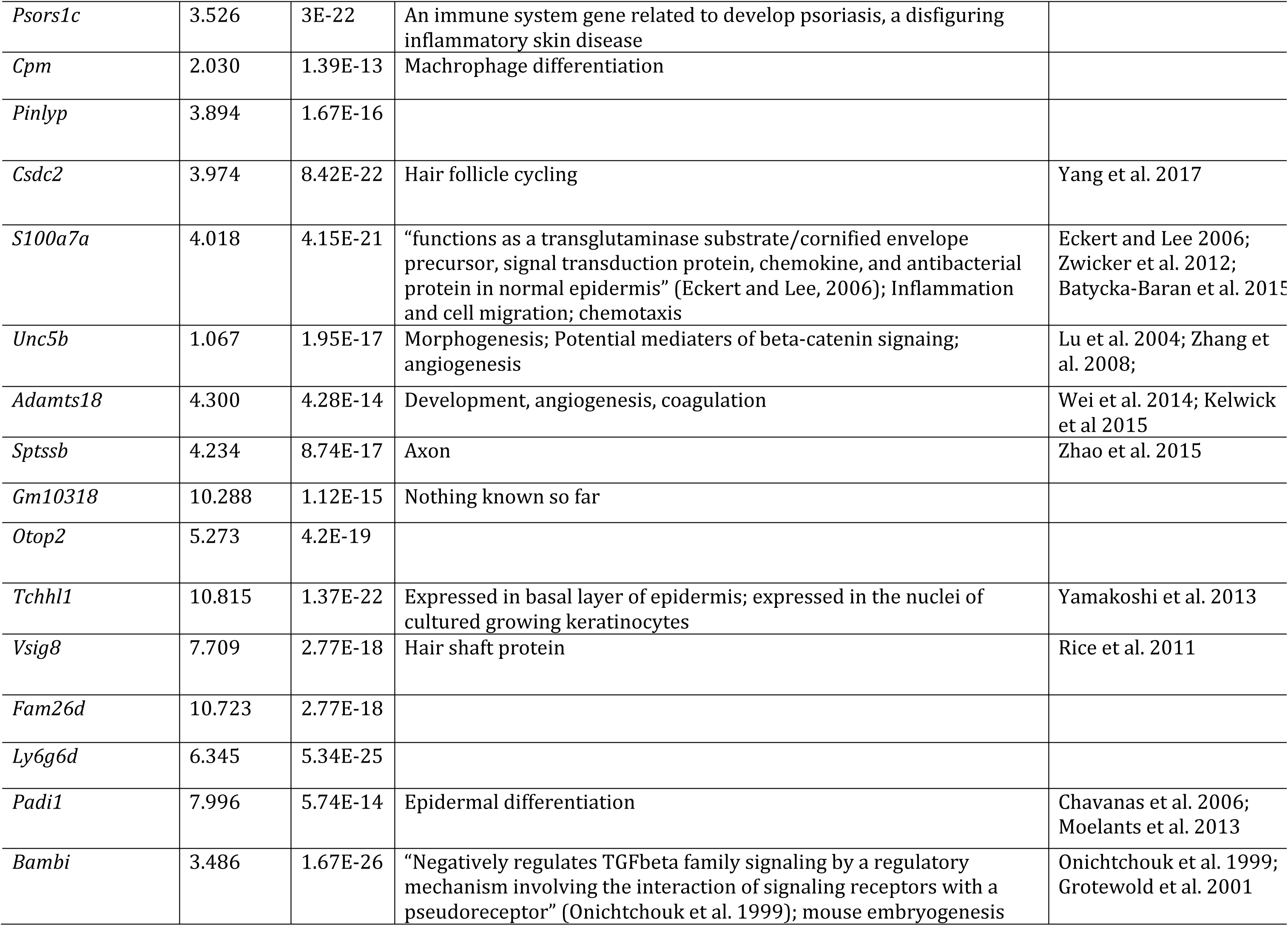
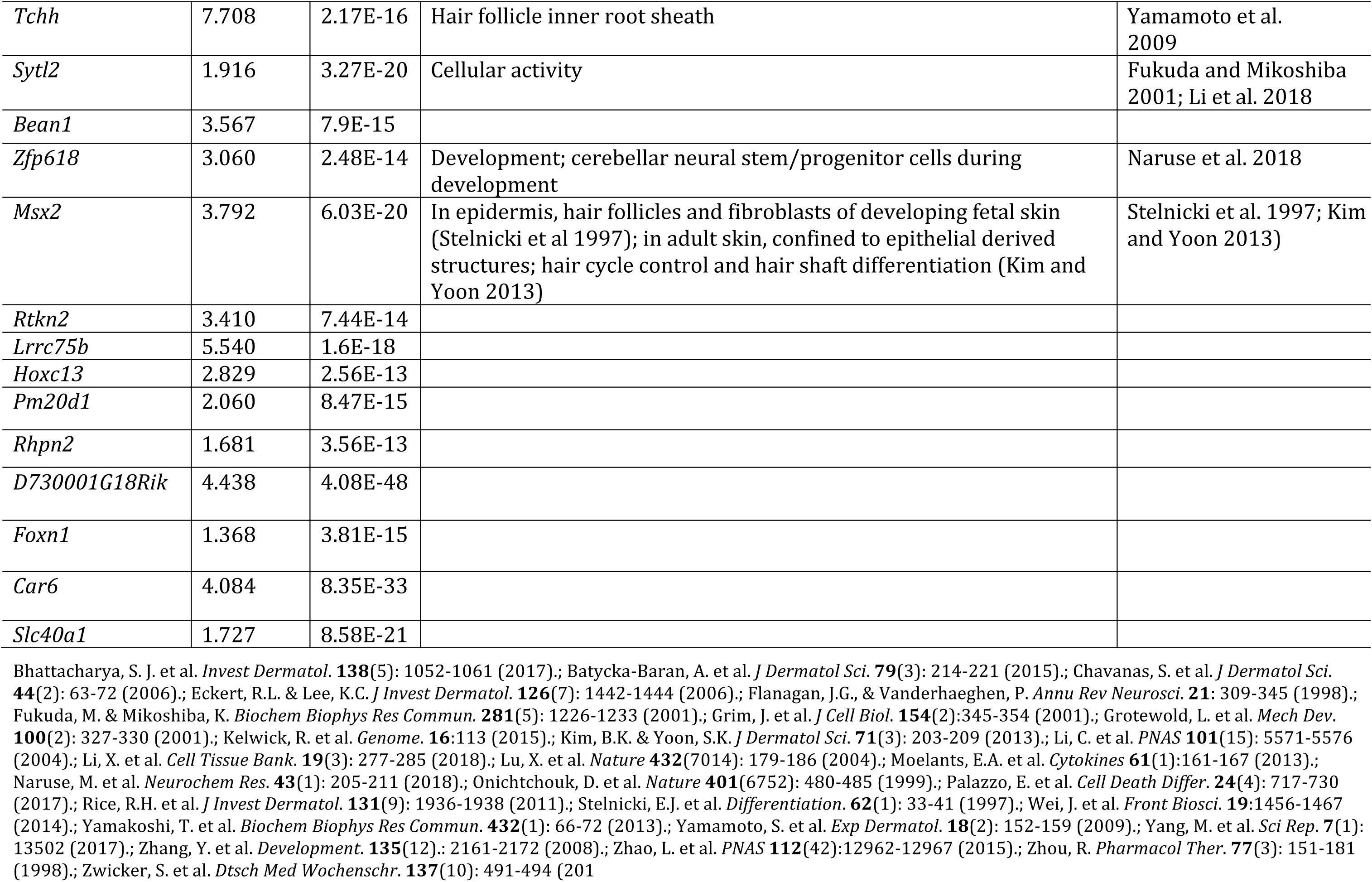
Top 50 genes altered in mice treated with oil and BCP. Top 50 genes were all up-regulated in the BCP group compared to the Oil group.

The transcriptome analysis prompted us to examine whether BCP treatment affected genes known as hair follicle stem cell markers [38]. The hair follicle bulge markers *Gli1*, *Lgr5*, and *Sox9* were significantly up-regulated (Log2FC, 2.4605, 1.236, 0.5377, respectively) in the BCP group compared to the Oil group (see bottom table in Fig. 8). Hair follicle infundibulum marker *Lrig1* was also significantly up-regulated by 40% (Fig. 8, log2FC 0.4035). These results on up-regulation of hair follicle bulge markers suggest that BCP treatment can stimulate hair follicle stem cell production.

**Fig 8.**
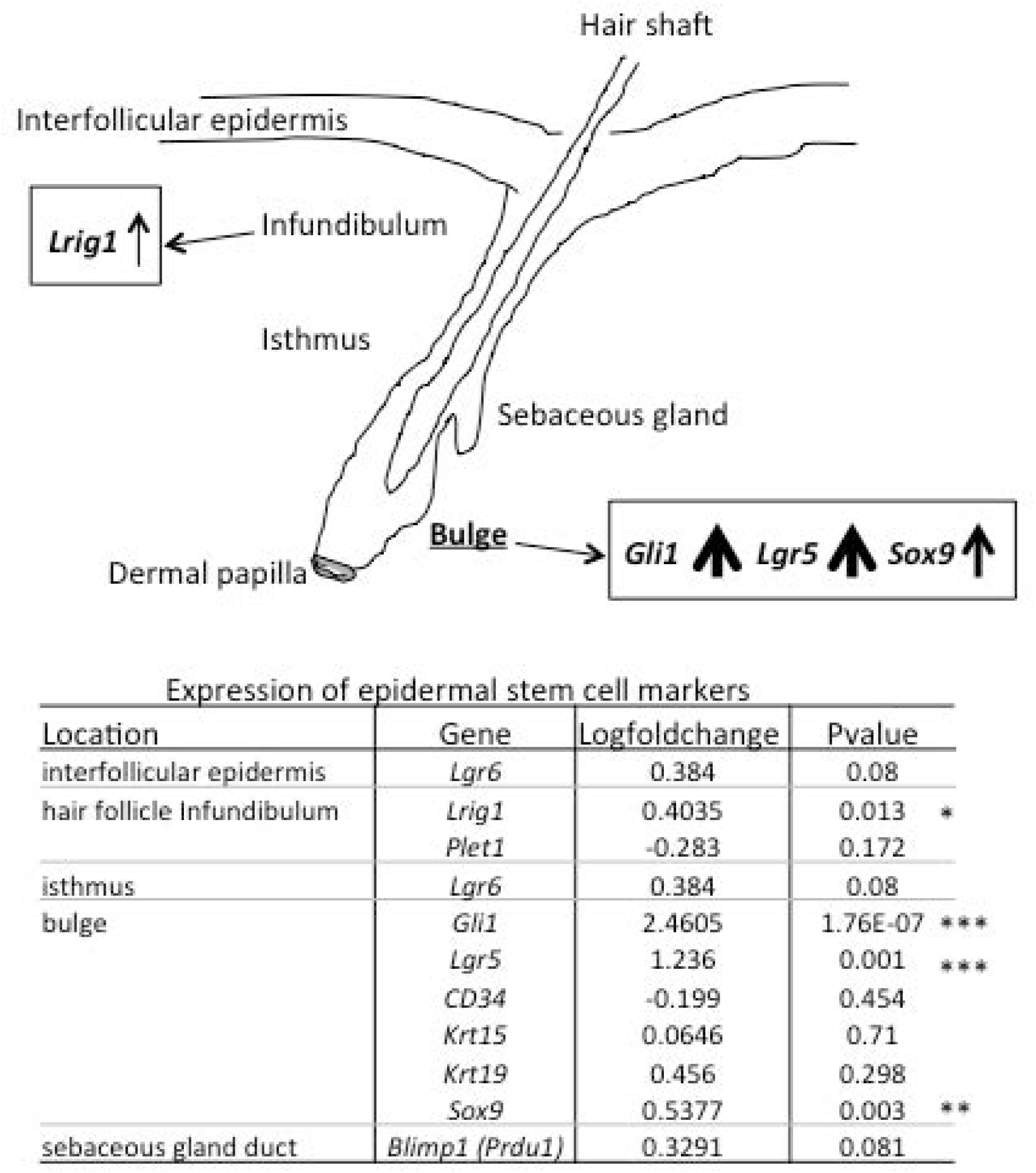
Comparison of the expression of hair follicle marker genes. Hair follicle genes related to the bulge were significantly up-regulated in the mice exposed to BCP compared to the Oil group. P-Values show the results of comparison between the BCP and the Oil groups. *: P<0.05, **: P<0.01, ***: P<0.001.

The inflammation stage of wound healing starts immediately after injury and pro-inflammatory cytokine levels usually peak within the first 10 to 20 hours [53, 54], then decrease for 2 days. The main source of cytokines is known to be neutrophils [53]. In our study, we used 17 to 18 hours post-injury skin and, of the pro-inflammatory cytokines, IL-1β and IL6 genes were significantly down-regulated in the BCP group compared to the Oil group (Table 2). This suggests that BCP treatment suppressed acute inflammation after skin exicision (Table 2). Studies have shown that there is a rebound in the inflammatory cytokines after 3 days, when fibroblasts migrate and new granulation tissues are formed, which suggest that the cytokines during the ‘rebound’ stage may play a role in wound *remodeling* [53]. Using the tissue harvested on post-surgery day 4, we examined the expression of the pro-inflammatory cytokines IL-1β and TNFα (Fig. 9a and b). IL-1β expression was especially strong in the wound bed of the BCP-treated mice. IL-1β and TNFα were expressed 2.7 times higher and 2.0 times higher, respectively, at the wound bed in the BCP group when compared to the Oil group (compared using color intensity in the image field). Apoptosis is involved in various stages of tissue repair with its main role in eliminating unwanted cells. If the acute inflammation immediately after injury is suppressed, it is possible to expect that the apoptosis rate might be suppressed in the BCP group. We conducted a terminal deoxynucleotidyl transferase (TdT) dUTP nick-end labeling (TUNEL) assay to detect apoptotic cells. TUNEL+ cells were abundant in the wound bed of both BCP-treated and oil-treated wounds when compared to the intact area. However, TUNEL staining was much stronger in the Oil-treated group (2.0 times higher in color intensity in the Oil group compared to the BCP group). In the BCP-treated wounds, TUNEL+ cells were present in the bottom of the epidermis and dermis close to the wound edge (Fig. 9c). These results suggest that, in the BCP group, pro-inflammatory cytokines are suppressed immediately following injury which may have produced reduced apoptosis at 4 days post-injury. Although the expression of pro-inflammatory cytokines is suppressed in the BCP group during the early stage after injury, they become more expressed than the Oil group at post-surgery day 4, which is when they are at the cell proliferation stage. Further studies are required to precisely determine the roles of these cytokines in the differences observed in wound healing between the BCP and Oil treated wounds.

**Fig. 9.**
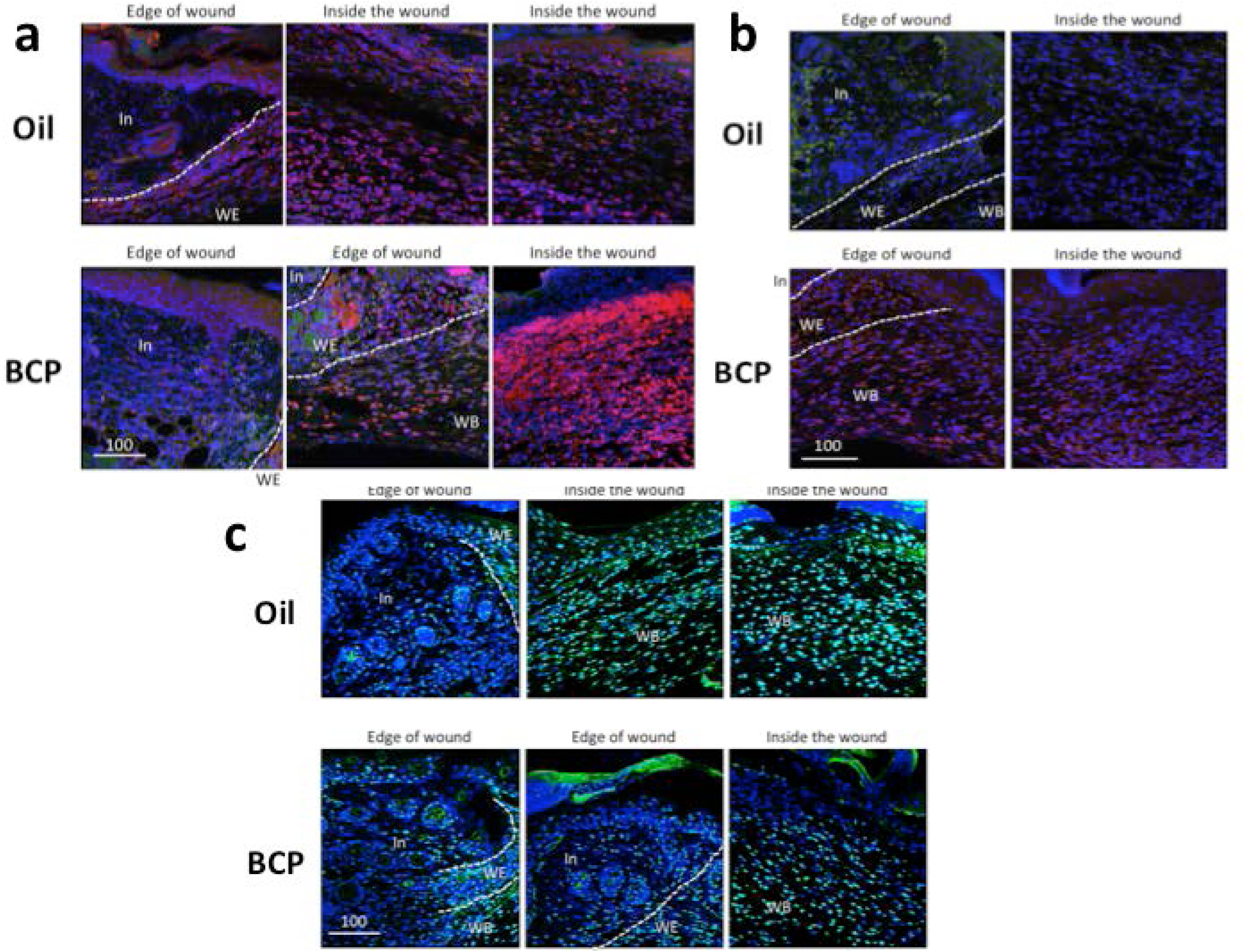
IL-1β, TNFα in the skin at the wound bed and close to the wound, and TUNEL staining of the skin at the wound bed and close to the wound. **(a)** IL-1β (red and purple) expression in the BCP and the Oil group. Blue color as well as purple color is Draq 5+ cells. Green is autofluorescence. **(b)** TNFα expression (red and purple) expression in the BCP and Oil group. Blue color as well as purple color is Draq 5+ cells. Green is autofluorescence. **(c)** TUNEL staining with double staining with Draq 5. Green and light blue is TUNEL+ cells. Blue and light blue is Draq5+ cells. Scale bars are μm.

**Table 2.**
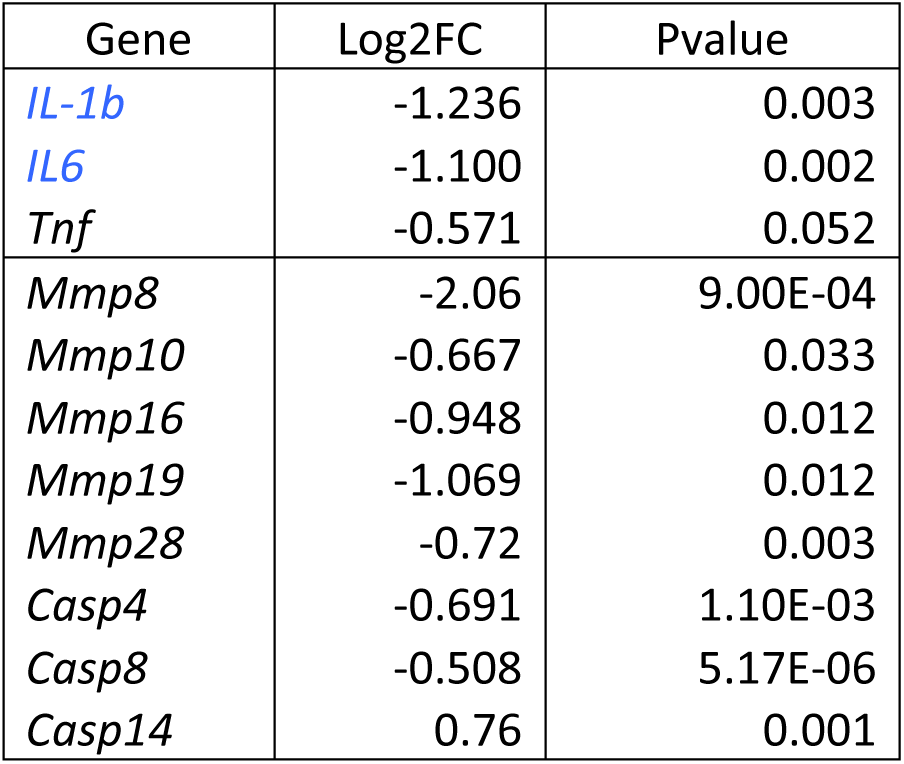
Expression of pro-inflammatory cytokines in the BCP group compared to the Oil group.

CB1 gene expression was not significantly different between the BCP and the Oil group, whereas CB2 gene was, to our surprise, down-regulated in the BCP group in the 17∼18 hours post-injury skin. Recent studies have shown that Transient Receptor Potential (TRP) channels become activated by some phytochemicals [55]. We examined the expression of genes related to TRP channels and found that TRPM1, TRPM6, TRPV4, and TRPV6 were significantly up-regulated and TRPM2 and TRPM3 were significantly down-regulated in the BCP group compared to the Oil group (Table 3). Importantly, there are growing number of papers showing the roles of TRP channels in the initiation of pain and itch perception as well as epidermal homeostasis and hair follicle regulation, suggesting that the TRP channels function as ‘ionotropic cannabinoid receptors’ [56]. Of the TRP channels designated as putative ionotropic cannabinoid receptors [56], the one that overlaps with TRP channels showing significant change after exposure to BCP was TRPV4, which showed significant up-regulation. TRPV4 is expressed in various cell types including sensory neurons and is considered to be a mechano/osmosensitive channel [57]. The results of our study and recent studies on the roles of TRP channels suggest that TRP channels could be involved in enhanced re-epithelialization by BCP.

**Table 3.**
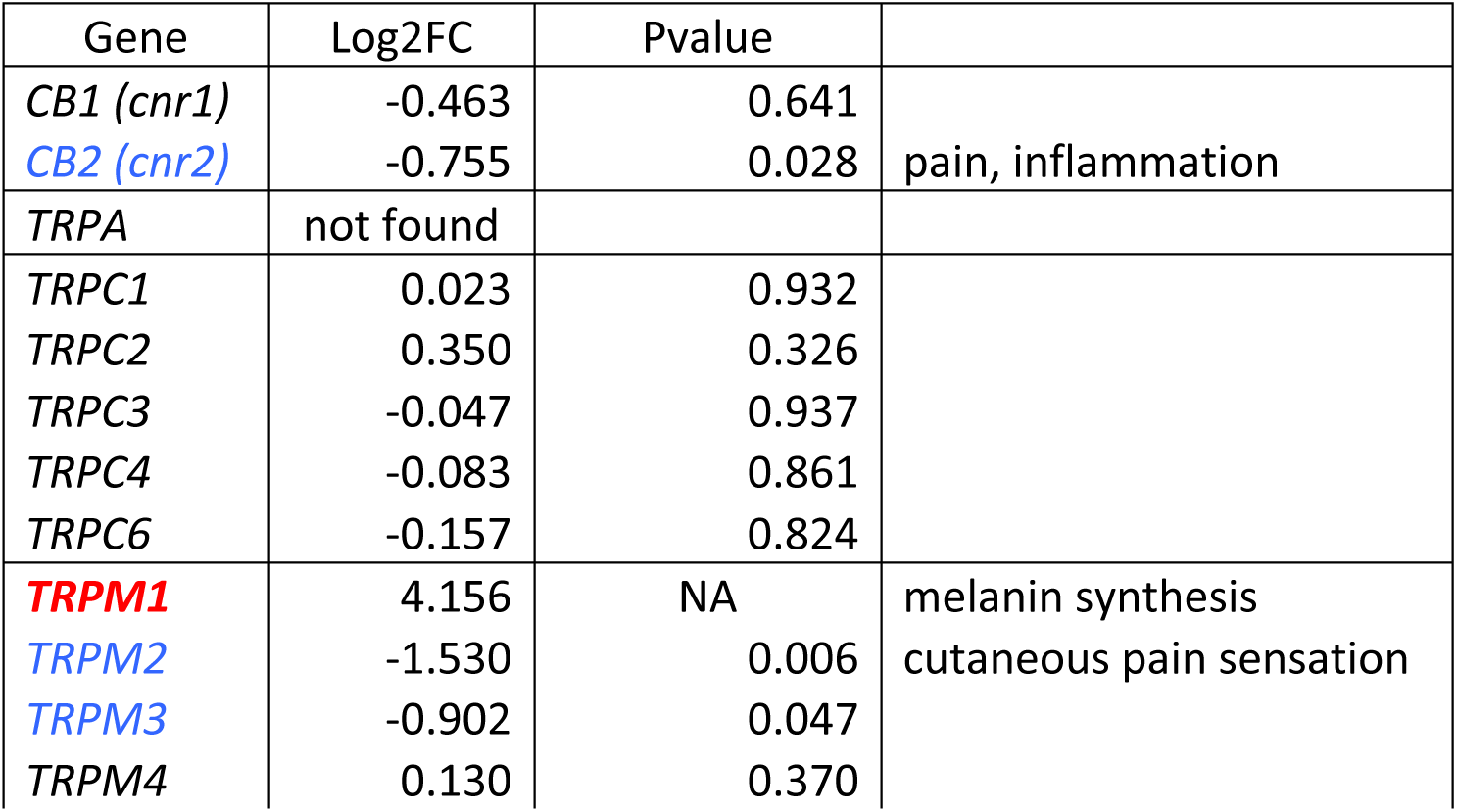
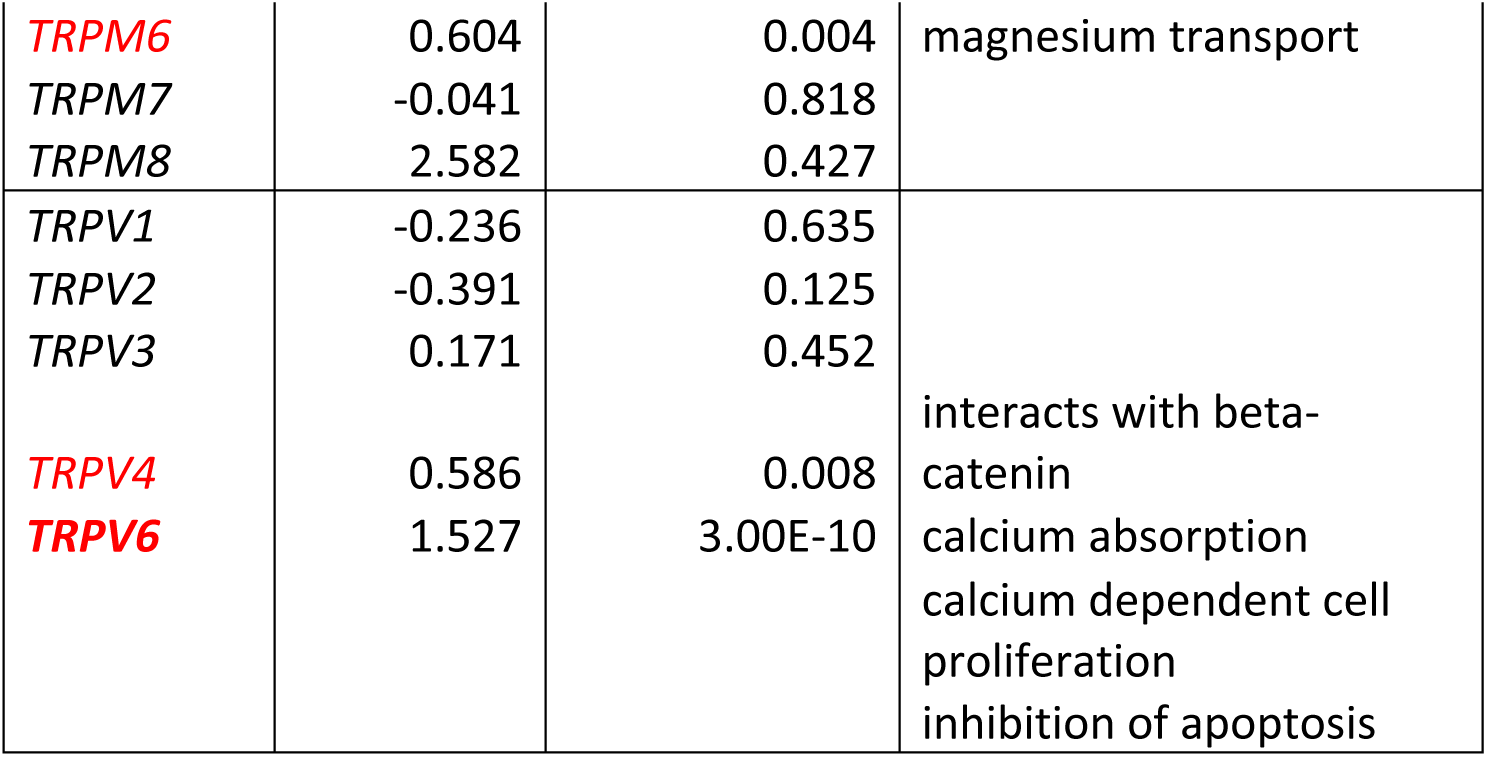
Expression of cannabinoid receptors and TRP channel genes. Genes up-regulated in the BCP group compared to the Oil group are written in red color and down-regulated genes are written in blue color. Genes written in black are not significantly different between the BCP and the Oil group. Genes written in bold type have P values over E-5. NA in P-Value is an outlier compared to other genes presumably because of its extremely high expression in the BCP group compared to the Oil group.

Ingenuity pathway analysis^®^ (IPA^®^) was performed using the gene expression results. In the comparison of the BCP vs. Oil group, signaling pathways related to inflammation and the immune system (TREM1 signaling) [58]were suppressed in the BCP group (S6 Fig.). In addition, genes that encode functions for cell proliferation and migration (e.g. the sonic hedgehog pathway, the planar cell polarity, the signaling pathway, fibroblast growth factor signaling pathway, and Wnt beta-catenin signaling pathway [S7 Fig. a, b, c, d]) were activated compared to the Oil group. Table 4 summarizes the genes in these signaling pathways that became significantly up/down-regulated in the BCP group compared to the Oil group. In each of these pathways, there were genes whose expression was dramatically increased (bold in Table 4). These could be the core genes affected by BCP: hedgehog interacting protein gene (*hhip*), sonic hedgehog gene (*shh*), and dual specificity protein phosphatase CDC14a (*cdc14a*) in the sonic hedgehog signaling pathway [59], frizzled-5 (*fzd5*), wingless-type MMTV integration site family member 5a, 10b, 11 (*wnt5a, 10b, 11*) in the planar cell polarity signaling pathway [60], fibroblast growth factor 22 and 23 (*fgf22, fgf23*) and phosphoinositide-3-kinase regulatory subunit 3 (*pik3r3*) in the fibroblast growth factor signaling pathway [61], and *wnt5a, 10b, 11*, whitefly-induced tomato gene (*wfi1*), transcription factor sox4 (*sox4*), and lymphoid enhancer-binding factor 1 (*lef1*) in the Wnt/beta-catenin signaling pathway [62]. Studies have found that, in the case of adult skin, hedgehog signaling is involved in epidermal homeostasis, *i.e*., the hair cycle and regrowth of hair [63]. *GLI1* is expressed in the hair follicle bulge and, when sonic hedgehog secreted from sensory neurons surrounding the hair follicle bulge stimulates the Gli1+ cells in the upper region of the bulge, the Gli1+ cells convert to multipotent stem cells. These multipotent stem cells migrate to epidermis and contribute to wound healing by becoming epidermal stem cells [64]. The planar cell polarity pathway is known to be involved in collective and directed cell movements during skin wound repair [65-68] and coordinating the directions of cell migration [67]. Studies using transgenic mice have shown that the role of fibroblast growth factor signaling includes keratinocyte proliferation [68]. Overall, the results on gene expression in our study suggest that exposure to BCP activated the pathways in a way to enhance 1) conversion of hair follicle bulge stem cells to epidermal stem cells, 2) keratinocyte cell proliferation, 3) coordinated and directed cell movement, which may have contributed to enhanced re-epithelialization by exposure to BCP.

**Table 4.**
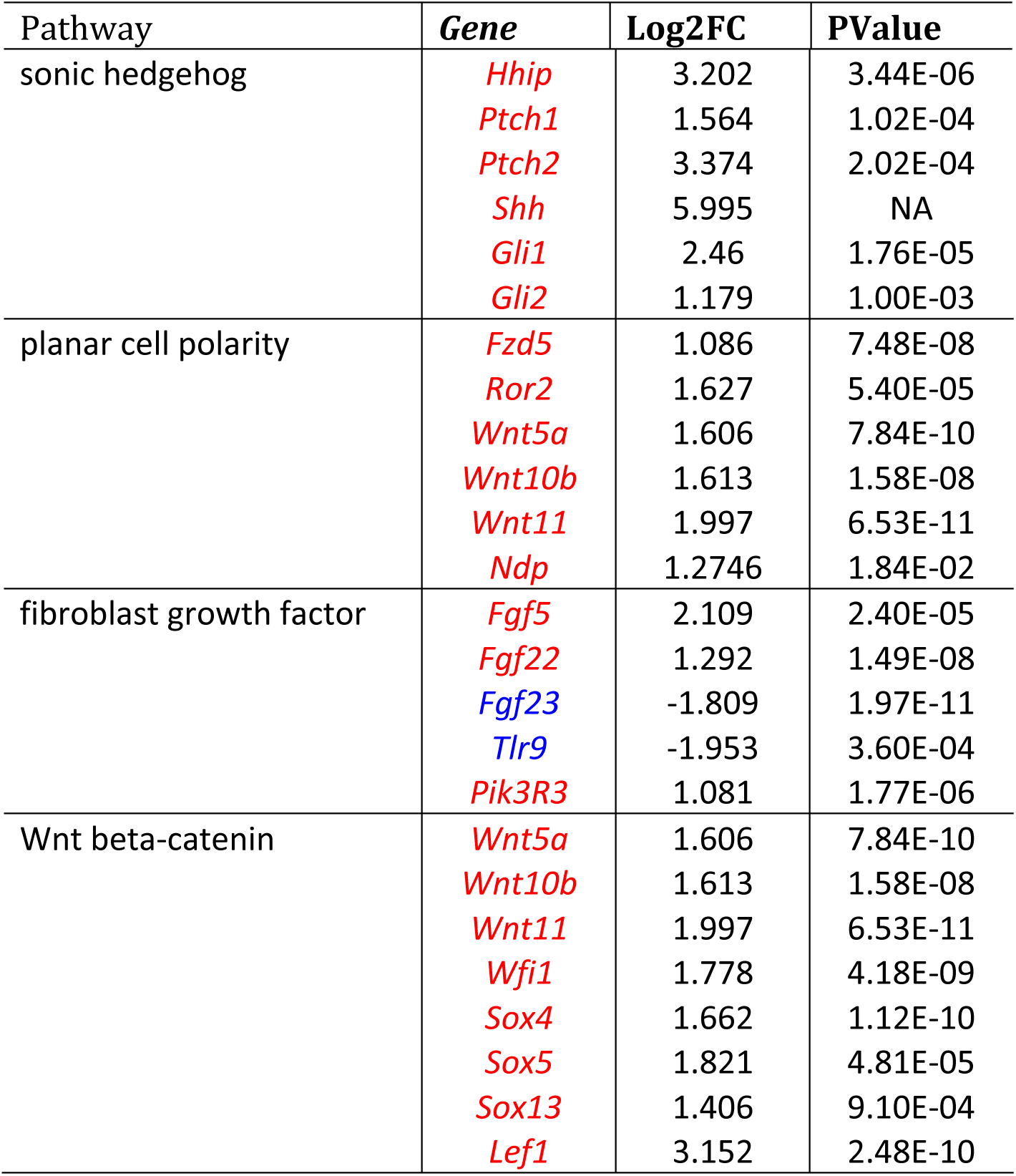
Significantly up/down-regulated genes in shh, pcp, fgf, and wnt/beta-catenin signaling pathways after exposure to BCP. Genes with log fold change over 1 are shown. Up-regulated genes are written in red, and down-regulated genes are written in blue. Results of genes with over E-05 P-values are shown in bold letters. NA in P-Values is an outlier compared to other genes presumably because of its extremely high expression in the BCP group compared to the Oil group.

### Sex differences in the impact of BCP

These results summarized above are based on experiments using female mice. Previous studies have shown sex differences in the morphology and physiology of mouse skin, with female mice having a 40 % thicker epidermis than males [69]. In addition, androgen receptors have an inhibitory influence on wound healing in males [70], which may hinder the impact of BCP.

We tested whether the influences of BCP on wound healing in male mice is similar to its influences in females and found no statistically significant differences in re-epithelialization between the BCP and Oil groups in males (based on K14 staining) (Fig 10). Whether males need a higher concentration of BCP to induce its impact or whether BCP does not affect males at all needs to be addressed in future.

**Fig 10.**
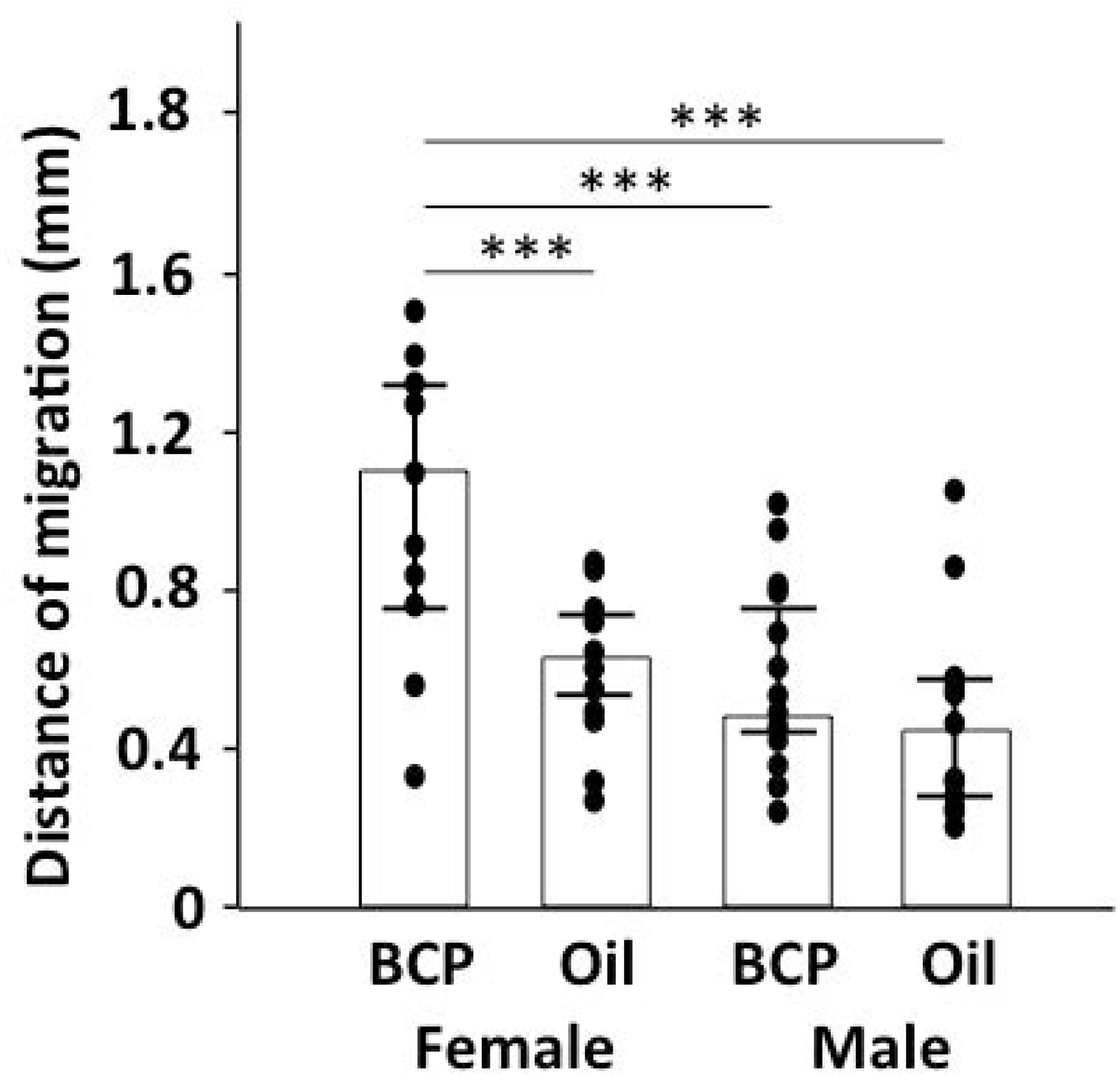
Influence of BCP on re-epithelialization in male and female mice. Length of K14+ staining area from the boundary of the intact and wounded area to the center of the wound in skin harvested on post-surgery day 4 from mice exposed to BCP or oil. Bar indicates median and line indicates 25 and 75% quartile. ANOVA, BCP vs. oil, *F*_1,66_=14.793, *P*<0.001; gender, *F*_1,66_=30.761, Tukey’s post-hoc test, ***: P<0.001; females, BCP, n=16, oil, n=16, males, BCP, n=23, oil, n=15 (76 words)

Sex steroids are known to affect wound healing [71], androgen receptors inhibit wound healing [60] whereas estrogen receptors accelerate it [32]. The re-epithelialization in the Oil control group for males was similar to that of the Oil group of female mice (Fig 10), which suggests that sex hormones did not suppress re-epithelialization in males.

### Olfactory receptors are not involved in wound healing

BCP is an odorant. When mice are topically treated with BCP, they are exposed to its smell and there is a possibility that the olfactory system has some role in the BCP mediated enhanced re-epithelialization. In addition, olfactory receptors are expressed in non-olfactory tissues. A synthetic analogue of sandalwood odorant, sandalore, is the ligand of the olfactory receptor gene OR2AT4 and that OR2AT4 receptors are expressed in human skin [72]. Thus odorants can directly produce an impact on non-olfactory tissues like skin tissues [72] other than through the route of the olfactory system to the brain, affecting emotional status [73]. There is a possibility that the impact of BCP was mediated by olfactory receptors expressed in the skin.

To examine if the olfactory system and/or olfactory receptors expressed in the skin has some role in enhanced re-epithelialization by exposure to BCP, we conducted multiple experiments. First we tested if mice can smell BCP by testing the expression of an immediate early gene protein in the olfactory bulb of mice exposed to BCP and found clear expression of c-FOS protein, suggesting that it is highly likely that mice can smell BCP (S8 Fig.). Then we tested whether exposure of injured mice to BCP only through the air would produce a similar impact as topical application, and found no improved re-epithelialization (S9 Fig). It was clear that the olfactory system was not significantly involved in the impact of BCP on re-epithelialization.

To examine if olfactory receptors expressed in the non-olfactory system are involved, we first examined the olfactory epithelium after exposure of non-injured mice to BCP for one hour to identify the olfactory receptor genes for BCP using pS6-IP RNAseq [74]. We could narrow down the candidates of olfactory receptor genes for BCP (S2 Table). It is known that odorants stimulate a limited number of types of olfactory receptors [75, 76]. As the BCP we used was not 100 % in purity, multiple olfactory receptor genes were up-regulated following exposure to BCP with *Olfr340* showing the largest fold change and significant P-Value (S2 Table). After this step, we analyzed the results of RNA sequencing of the skin to determine if the olfactory receptor genes up-regulated in olfactory epithelium by exposure to BCP are expressed in skin and are up-regulated after topical application of BCP. There were many olfactory receptor genes expressed in the skin (S3 Table), and one of them overlapped with the olfactory receptor genes up-regulated in olfactory epithelium (*Olfr111*), but none of them were up-regulated specifically in the BCP group. From these results, we conclude that olfactory receptors expressed in skin are not involved in the enhanced re-epithelialization by exposure to BCP.

### Synergetic influences by exposure to BCP

Wound healing starts from the inflammatory stage and shifts to the cell proliferation stage. Since BCP is an agonist of CB2, we had assumed that it can improve wound healing by decreasing pain and reducing pro-inflammatory signals and speed up the shift to the cell proliferation stage. However, our results suggests that BCP may directly impact cell proliferation and cell migration concomittent with reduced inflammation. BCP also stimulates mitogenesis of hair follicle stem cells. Hair follicle stem cells participate in re-epithelialization when there is a cutaneous wound [41, 51], our study suggests that enhanced re-epithelialization by exposure to BCP is mediated by BCP stimulated production of hair follicle stem cells in the bulge, and their migration to epidermis and to wound bed.

BCP is known to suppress apoptosis [77] and necroptosis [78], raising the possibility that suppression of cell death enabled higher cells survival and proliferation. The suppression of *Septin4* (*Sept4*) has been shown to protect cells from cell death and promote wound healing [79], however, *Sept4* was not up-regulated in our study and seemed to be not involved in BCP enhanced re-epithelialization. It is possible that exposure to BCP generates a specific wound healing environment by up-regulating the genes related to embryonic growth as was suggested from our results on transcriptome analyses.

Our study focused on the early stage following skin excision. Studies have shown that oxidants from inflammatory cells are involved in fibrotic healing (scarring) [80, 81], which suggests that suppression of inflammation through activation of CB2 by BCP, in addition to up-regulation of genes related to embryonic growth would have a high possibility of reducing scar formation. These possibilities need to be addressed in future by testing the influence of exposure to BCP on the late stage after injury.

### Odors affect us more than we think

It has been several decades since the first report that olfactory receptors are expressed in non-olfactory regions [82] and that odors have functions other than olfactory and pheromonal communication [83]. Our observations that BCP activates multiple types of receptors in the skin further unveils the complexity of chemical signaling. Although we humans typically rely on visual and auditory stimuli in communication, it is important to understand how odors in the environment, whether it is from other individuals [1] or from plants [2-6], could be affecting our condition. In addition, BCP is but one component commonly found in essential oils of herbal extracts. Further examination of the compounds in herbal extracts could lead to additional useful molecules that could benefit our health.

## Materials and methods

### Mice strains

We used a standard murine model for wound healing. Mice were used between 8 to 10 weeks in age in *in vivo* experiments. C57BL/6J, and CB2-/- mice were used in experiments. Primary cells for *in vitro* experiments were obtained from P0 to P1 neonate pups or adult mice from C57BL/6J, or CB2-/- mice. In *in vivo* experiments, C57BL/6J mice were used. All mice were kept on a 14:10, L/D light cycle, and water and food were provided *ad libitum*. All the procedures were following a protocol (#16-022) approved by the Indiana University Institutional Animal Care and Use Committee (IACUC).

### Surgery

We studied small full-thickness excisions on the backs of adult mice and developed methods to hold the treatment buffer solution on the wounded area (S1 Fig a). Hair of the dorsal area was removed and the area for surgery was cleansed using betadine and 70% ethanol. A small piece of the dorsal skin of a mouse was excised (< 5mm x 5mm) under anesthesia. A silicone ring was attached to the wounded area using a liquid sealing bandage (S1 Fig a center) (OD: 15mm, ID: 8mm, thickness, 3mm). This ring served as a reservoir and 50 µL of BCP (50 mg/kgbw; 0.2nM) (W225207, Sigma-Aldrich, Co. St. Louis, MO) diluted in olive oil, or olive oil as control (O1514, Sigma-Aldrich, Co. St. Louis, MO), was placed in the reservoir. The gas chromatography-mass spectrometry (GC/MS) analysis of the BCP standard showed that it contained 81.8% BCP and 14.0% α-caryophyllene (α-humulene). The remaining 4.2% consisted of 1.6% caryophyllene oxide and minor amounts of other molecular weight 204 sesquiterpenes (0.1-0.8% each) (S4 Fig., S1 Table). Molecular weights were calculated by the amount of BCP (81.9% of the volume converted to molecular weight).

The ring was covered with a baked PDMS lid (polydimethylsiloxane) (Dow Corning Sylgard^®^ 184 Silicone Elastomer Kit; Dow Corning, Auburn MI) and sealed with liquid bandage to hold the BCP media or olive oil. A gauze bandage was used to loosely cover the body to prevent the mouse from removing the ring with PDMS lid (S1 Fig 1a, right down). The bandage and buffer media were removed daily and replaced with a new bandage and buffer media. The wounded area was collected on the 3^rd^, 4^th^, and 5^th^ day, and immediately fixed with 10% formalin for one day, then soaked in 70% ethanol.

### Gas chromatography – mass spectrometry (GC/MS) analysis of BCP

BCP was diluted to 180 ng/µL with a tert-butyl methyl ether solvent (99+%, Aldrich Chemical Company, Milwaukee, WI). The BCP composition was analyzed using 2 µL of the diluted BCP in a thermal desorption tube of the Thermal Desorption Autosampler and Cooled Injection System (TDSA-CIS 4 from Gerstel GmbH, Mülheim an der Ruhr, Germany) connected to an Agilent6890N Gas chromatograph – 5973iMSD Mass spectrometer (Agilent Technologies, Inc., Wilmington, DE). BCP components were identified with spectra through the NIST Mass Spectral Search Program for the NIST/EPA/NIH Mass Spectral Library (Version 2.0 a, 2002). Additionally, the mass spectra and Kovats retention index matches were verified with the reference spectra from Adam (2001) [84]. Caryophyllene oxide was verified with the standard (99%) from Sigma-Aldrich (St. Louis, MO).

### CB2 agonist and antagonist treatment

CB2 agonist JWH133 (1 µM; Cat. 10005428, Cayman Chemical Co., Ann Arbor, MI), dissolved in alcohol and kept at −20°C, was dissolved in olive oil after the alcohol was evaporated and used instead of BCP in the daily treatment on the wounded area. CB2 antagonist, AM630 (4 mg/kgbw (7.9 µM/kgbw); SML0327, Sigma-Aldrich Corp. St. Louis, MO) was dissolved in saline and injected 20 min before the daily treatment of BCP.

### Immunofluorescence staining to evaluate re-epithelialization

Mice were euthanized by cervical dislocation and the wounded areas were harvested, fixed in 10% formalin at room temperature and then in 70% ethanol at 4°C until they were embedded in paraffin. Samples were sectioned at 10 µm, deparaffinized and hydrated, went through antigen retrieval using the sodium citrate buffer (10mM, >95°C for 20 to 30 min), blocked with 10% NGS, and primary stained (see below for antibodies) overnight at 4°C. Then sections were washed, secondary stained, nuclear stained using Draq 5, washed (3 times), and covered with Prolong Diamond (ProLong™ Diamond Antifade Mountant P36965, Thermo Fisher Scientific).

Antibodies used are as follows: proliferating cell nuclear antigen (PCNA (PC10) mouse mAb #2586, Cell Signaling Technology, Danvers, MA, 1:1000), keratin 14 (K14, rabbit pAb, 905301, 1:1000, BioLegend, San Diego, CA), PDGFRalpha (mouse mAb, sc-398206, Santa Cruz Biotechnology, CA, 1:1000), vimentin (rabbit, 10366-1-AP, Protein Tech, 1:500), estrogen receptor alpha (Eralpha, mouse mAb, sc-51857, sc-8002, Santa Cruz Biotechnology, 1:250), filaggrin (rabbit pAb, BioLegend, San Diego, CA, 1:1000), TNFalpha (rabbit pAb, ab6671, Abcam, 1:100), IL-1beta (rabbit pAb, ab9722, Abcam, 1:100). Alexa fluor 594 anti-rabbit and Alexa fluor 488 anti-mouse (both Invitrogen, 1:500). Stained sections were observed using laser scanning confocal microscope Leica SP5 at the IUB Light Microscope Imaging Center.

### TUNEL staining

Mice were euthanized by cervical dislocation and the wounded area was harvested, fixed in 10% formalin at room temperature and then in 70% ethanol at 4°C until embedded in paraffin. Samples were sectioned at 10 µm, deparaffinized and hydrated. TUNEL staining was conducted following manufacturer’s protocol (In Situ Cell Death Detection Kit, Version 17, Roche Diagnistics GmbH, Mannheim, Germany). Briefly, slide glasses were first rinsed with PBS for 5 min twice, then went through antigen retrieval using sodium citrate buffer (10mM, >95°C for 20 to 30 min), and rinsed with PBS for 10 min three times. Blocked with 0.1 M Tris-HCl, pH7.5 with 3% BSA and 20% normal bovine serum for 30 min. Slides were rinsed twice with PBS for 10 min twice. Then TUNEL reaction mixture were added and slides were incubated for 1 hour at 37°C. Slides were rinsed for 10 min once, stained with Draq5 for 10 min at room temperature, and rinsed for 10 min three times. Finally, slides were covered with Prolong Diamond (ProLong™ Diamond Antifade Mountant P36965, Thermo Fisher Scientific). Stained sections were observed using laser scanning confocal microscope Leica SP5 at the IUB Light Microscope Imaging Center.

### BrdU injection and staining

On post-surgery day 4, mice were injected with bromodeoxyuridine (BrdU) (300 mg/kgbw) twice every 2 hours, and the skins were harvested 2 hours after the second injection. Harvested skin was fixed with 10% formalin, and then with 70% ethanol on the following day. Samples were embedded in paraffin and cut into sections using microtome at 10 µm. Sections were then deparaffinized and hydrated, went through antigen retrieval using 4N HCl at 37°C for 30 min, washed with 0.1M PBS, blocked with 10% NGS, and went through overnight staining at 4°C with anti-BrdU (1:300, rat, Accurate Chemical Co. #OBT0030). On the second day, sections were washed with 0.1M PBS, stained with anti-rat Alexa fluor 488 (1:500, Invitrogen) at room temperature, washed, stained with Draq5, washed, and covered after placing one drop of Prolong Diamond (ProLong™ Diamond Antifade Mountant P36965, Thermo Fisher Scientific). Sections were observed using laser scanning confocal microscope (Leica DMI 6000 CS inverted microscope) in the IUB Light Microscope Imaging Center and analyzed using Fiji/ImageJ software.

### Immediate Early Gene (*c-fos*) protein expression experiment and staining

Each mouse was exposed to BCP or male murine pheromone 2-sec-butyl-4,5-dihydrothiazole (SBT) (250ppm, positive control) [85, 86] for 1 hour in a clean cage. Then mice were anesthetized with 2.5% isoflurane and cardiac perfused with saline for 2 min, followed by 4% paraformaldehyde for 6 to 7 min. The brains were dissected out and post-fixed with 4% paraformaldehyde overnight at 4°C, transferred to PBS with 15% sucrose and kept at 4°C. The brains were embedded in OCT on dry ice and cryosectioned using Leica CM1850 at 20 µm and kept at −20°C until use.

For the staining of c-*fos* gene protein, the brain sections were washed 3 times for 10 mins with 0.1M PBS, blocked with 10% NGS, 0.5% Triton X-100 in 0.1M PBS for 1 to 2 hours, and went through primary staining with anti-cFOS (Invitrogen OSR00004w, rabbit, 1:250; Abcam ab190289, rabbit, 1:2000) overnight at 4°C. Then sections were washed 3 times for 10 mins and went through secondary staining (Alexafluor 594, anti-rabbit, 1:500), washed with 0.1M PBS, stained with Draq5, washed (0.1M PBS 3 times each 10 mins), and covered with Prolong Diamond. Bregma distance about 4.28mm was used in the comparison. The stained sections were imaged using laser scanning confocal microscope Leica SP5 at the IUB Light Microscope Imaging Center. The numbers of cfos+ cells in the whole glomerular layer, external plexiform layer, mitral layer, internal plexiform layer, and granule cell layer of olfactory bulbs were counted using Fiji/Image J.

### Isolation of primary cells

#### Fibroblast isolation

Mice (neonate or adult) were euthanized by cervical dislocation and cleansed using betadine, sterlized water, and 70% ethanol. The harvested skins were treated with 0.25% trypsin (Gibco, 15050065, Waltham MA) at 4°C overnight. Dermis was collected after trypsin treatment, minced and pipetted to dissociate, and incubated at 37°C with collagenase A buffer in cell culture media (1 mg/mL). Following the incubation, the buffer with dissociated cells was filtered using 100 µL strainer, and centrifuged at 500 X G for 8 min. Supernatant was removed and the pellet was re-suspended in CnT-PR-F media (CELLnTECH; Zen-Bio, Research Triangle Park, NC), filtered with 40 µm strainer, and centrifuged again at 500 X G for 8 min. Following another re-suspension to new culture media, cells were seeded on petri dishes to culture at 37°C with 5% CO_2_.

#### Keratinocytes

Skins were collected from neonate mice after the mice were decapitated and cleansed. Collected skins were placed in petri dish with 0.25% Trypsin (Gibco, 15050065, Waltham MA), with their epidermis side up and above the Trypsin buffer, at 4°C for 6 to 8 hours. Then the epidermis of the skin was removed from dermis, minced and pipetted to dissociate in the trypsin buffer, filtered using 70 or 100 µm strainers, and centrifuged at 500 X G or 800G for 8 min. Supernatant was removed and the pellet was re-suspended in high calcium DMEM medium ((Gibco, 10569-010, Waltham MA) supplemented with FBS (10%), p/s, and Glutamax), filtered using 40 µm, centrifuged again and resuspended with the high calcium DMEM medium. Then cells were seeded on Collagenase I coated petri dishes (Collagen I, Coated Plate 6 Well, Gibco Life technologies A11428-01, Waltham MA), incubated at 32 to 35°C^45^ with 5% CO_2_ with high calcium DMEM medium overnight^46^. On the following day, culture medium was replaced with low calcium medium (CnT-PR, CELLnTECH; Zen-Bio, Research Triangle Park, NC) and penicillin streptomycin (1140122, ThermoFisher Scientific, Waltham MA).

### Chemotaxis assay and scratch tests

#### Chemotaxis assay

Primary cultured cells (300 µL) were seeded in the inserts of 24 well cell migration plate (Cell Biolabs Inc, CBA100, CytoSelect 24 well cell migration assay kit) with 500 µL of DMEM cell culture media containing BCP (5 µL in 10 µL of DMSO in 10mL of cell culture media) or DMSO (control; 10 µL of DMSO in 10 mL of cell culture media) in the well, and incubated at 37°C and 5% CO_2_ for 1 day (approximately 18 to 24 hours). Then the inserts were washed with water and observed under microscope, photographed using EVOS XL Core (ThermoFisher Scientific, Waltham MA), and the number of the cells on the outside bottom of the insert was counted using Fiji/ImageJ.

#### Scratch tests

Primary cultured cells were seeded in 12 well plates. Two days later the culture media was changed to either 2 mL of DMEM cell culture media containing BCP (0.06 µL in 10 µL of DMSO in 10 mL of cell culture media) or DMSO (control; 10 µL of DMSO in 10 mL of cell culture media) and the surface of the bottom of the wells was scratched with the tip of 1mL pipet tips. Photos were taken immediately after the scratches were made, 6 hours later, and 1 day later. Widths of the scratch line were measured using Fiji/Image J.

### Time lapse *in vitro* observation of cell proliferation

Primary cultured cells were exposed to BCP in DMEM culture media or DMSO (control) in DMEM culture media on the next day after the cells were seeded on petri dishes with cover glass bottom (12-567-401, Thermo Scientific^TM^Nunc^TM^). Immediately after the media was changed to the media with BCP or the media with DMSO alone, cells were cultured on an Olympus OSR Spinning Disk Confocal microscope with Tokai Hit Stage Top Incubation system kept at 37°C and 5% CO_2_. DIC images were taken every 3 mins for 7 hours using a Hammamatsu Flash 4V2 camera and movies were generated using Metamorph software. Obtained images were played and the number of dividing cells were counted.

### Transcriptome analyses using skin

The wounded part of the skin was exposed to either BCP or olive oil (control) and the tissues were harvested 17 hours later. This was based on the study which reported the time course in the expression of genes related to inflammatory responses showing the peak within the first 20 hours [54]. Tissues from another group of mice were harvested similarly but without wound (no-injury control). The wounded area and its surrounding area (2 x 2 cm) were immediately dissected out and RNA extraction was conducted using Maxwell^®^ RSC simplyRNA Tissue Kit (AS1340, Promega, Madison WI) following the manufacturer’s protocol. The volume of extracted RNA was measured using Epoch™ Microplate Spectrophotometer (BioTek, Winooski, VT) with Take 3 Micro-Volume Plates (BioTek, Winooski, VT) and Gen5 Microplate Reader and Imager Software (BioTek, Winooski, VT).

The integrity was analyzed with Agilent Technologies 2200 Tape Station (Agilent Technologies). The input was quantified by using Qubit RNA BR Assay Kit and RNA-seq was conducted using NextSeq Series RNA-seq Solution (Illumina) at the Center of Genomics and Bioinformatics, IUB. Samples were prepared using TruSeq Stranded mRNA HT Sample Prep Kit (Illumina). Sequencing was performed using an Illumina NextSeq 500/550 Kits v2 reagents with 75 bp sequencing module generating 43 bp paired-end reads. After the sequencing run, demultiplexing was performed with bcl2fastq v2.20.0.422.

### pS6-IP RNA-Seq analyses using olfactory epithelium

To perform pS6-IP RNA-Seq, we used 4 pairs of litter-matched mice. We used both male and female mice in both control and experimental conditions with no specific orientation. Single pairs of mice were habituated in clean paper tubs in a fume hood for 1 hour. Following habituation, mice were transferred to another paper tub with an uni-cassette containing a 2 cm x 2 cm filter paper spotted with either water (control) or 1% (v/v) 10 L BCP dissolved in water (experimental) for 1 hour.

Following odor stimulation, mice were killed and the olfactory epithelium (OE) was dissected in 25 mL of dissection buffer (1× HBSS [Gibco, with Ca^2+^ and Mg^2+^], 2.5 mM HEPES [pH 7.4], 35 mM glucose, 100 μg/mL cycloheximide, 5 mM sodium fluoride, 1 mM sodium orthovanadate, 1 mM sodium pyrophosphate, 1 mM beta-glycerophosphate) on ice.

The dissected OE was transferred to 1.35 mL of homogenization buffer (150 mM KCl, 5 mM MgCl_2_, 10 mM HEPES [pH 7.4], 100 nM Calyculin A, 2 mM DTT, 100 U/mL RNasin (Promega), 100 μg/mL cycloheximide, 5 mM sodium fluoride, 1 mM sodium orthovanadate, 1 mM sodium pyrophosphate, 1 mM beta-glycerophosphate, protease inhibitor (Roche, 1 tablet per 10 ml)) and homogenized three times at 250 rpm and nine times at 750 rpm (Glas-Col). The homogenate was transferred to a 1.5 mL lobind tube (Eppendorf), and centrifuged at 2,000 X G for 10 min at 4°C. The supernatant was then transferred to a new 1.5 mL lobind tube, to which 90 μL 10% NP-40 (vol/vol) and 70 μL 300 mM DHPC (Avanti Polar Lipids) were added. The mixture was centrifuged at 17,000 X G for 10 min at 4°C. The supernatant was transferred to a new 1.5 mL lobind tube and mixed with 6 μL pS6 antibody (Cell Signaling, D68F8). Antibody binding was allowed by incubating the mixture for 1.5 hour at 4°C with rotation. During antibody binding, Protein A Dynabeads (Invitrogen, 100 μL per sample) was washed three times with 900 μL beads wash buffer 1 (150 mM KCl, 5 mM MgCl_2_, 10 mM HEPES [pH 7.4], 0.05% BSA (wt/vol), 1% NP-40). After antibody binding, the mixture was added to the washed beads and gently mixed, followed by incubation for 1 hour at 4°C with rotation. After incubation, the RNA-bound beads were washed four times with 700 μL beads wash buffer 2 (RNase-free water containing 350 mM KCl, 5 mM MgCl_2_, 10 mM HEPES [pH 7.4], 1% NP-40, 2 mM DTT, 100 U/mL recombinant RNasin (Promega), 100 μg/mL cycloheximide, 5 mM sodium fluoride, 1 mM sodium orthovanadate, 1 mM sodium pyrophosphate, 1 mM beta-glycerophosphate). During the final wash, beads were placed onto the magnet and moved to room temperature. After removing supernatant, RNA was eluted by mixing the beads with 350 μL RLT (Qiagen). The eluted RNA was purified using RNeasy Micro kit (Qiagen). Chemicals were purchased from Sigma if not specified otherwise.

cDNA from purified RNA were generated using SMART-Seq v4 Ultra Low Input RNA Kit (Takara Clontech). Libraries were made using the Nexterra XT DNA Library Preparation Kit. Up to 12 samples were pooled and sequenced on Illumina HiSeq2000/2500 on single-end read mode. RNA-Seq data was aligned using Kallisto. Differential expression analysis was performed using a combination of DESeq and EdgeR packages in R.

### Open-field behavior test

The behavior of the mice was observed on post-surgery days 1 and 3. Three groups of mice were observed, *i.e*., the no-injury control group, the Oil control group, and the BCP group. The no-injury control group had the dorsal hair removed and a bandage was placed on the day when other mice went through surgeries, but no incision was made. Bandages of these mice were changed daily as well. The olive oil control group and the BCP group underwent the procedure detailed above. A mouse was placed in a cup (about 10 cm diameter x 15 cm height) and placed in the center of an open field test apparatus (measurements: width 50 cm x depth 25 cm x height 30 cm) covered with the cup. Video recording using a camera (Logitech QuickCam Pro9000) and bTV software was conducted either from the top for analysis of the amount of traveling or from the side for analysis of behavior patterns. After camera was started, the cup covering the mouse was removed and behaviors were recorded for 5 minutes. The behaviors recorded from the top were analyzed using Ethovision (Version 7, Noldus, Leesburg, VA) and behaviors recorded from the side were analyzed using ELAN software (http://tla.mpi.nl/tools/tla-tools/elan/, Max Planck Institute for Psycholinguistics, The Language Archive, Nijmegen, The Netherlands) [87]. Behaviors were categorized based on the definition given by previous studies [88-90] and quantified using ELAN software. The behaviors coded using ELAN were then exported to Microsoft Excel where average time for each behavior, incidence of each behavior, and total time for each behavior were determined.

### Statistical analyses

ANOVA and Student T-tests were used for comparison of the groups. Tukey post-hoc test was used for pair-wise comparison. Comparison of % was conducted using Fisher’s exact test of indepence.

## Acknowledgements

This project was funded by the Office of the Vice Provost of Research at Indiana University Bloomington through the Collaborative Research and Creative Activity Funding Award to S.K. while she was Associate Research Professor at the Medical Science Program of the School of Medicine of IU. Large amount of this project was conducted while S.K. was Associate Research Professor at the Medical Science Program of the School of Medicine of IU, financially supported by the previous Director of Medical Science Program, Prof. John Watkins III. S.K. thanks Dr. A. Poehlman for her help in editing English. S.K thanks A. Straiker of the Department of Psychological and Brain Sciences, IU, for his kind gift of CB2 agonist and antagonist used in the project and also for the advice on CB2. S.K thanks J. Wager-Miller of the Department of Psychological and Brain Sciences, IU, for his kind gift of CB2 knockout mice in the project. S.K. thanks J. Foley of Medical Science Program of IU School of Medicine to let S.K. use part of his laboratory space and resources for the project. S.K. thanks S. Childless of Medical Science Program of IU School of Medicine for her help in sectioning samples and J.K. Leffel II of Department of Psychological and Brain Sciences, IU, for his help on using Ethovision. RNA sequencing and bioinformatic analyses was conducted at the Center for Genomics and Bioinformatics of IUB and S.K. thanks A.M. Buechlein, D.B. Rusch and J. Huang for their services. S.K. also thanks K. Mackie of the Department of Psychological and Brain Sciences for his advice on the project, J. Powers of the IU Light Microscopy Imaging Center (LMIC) for his help and advice while using LMIC, and D. Sinkiewicz of the Center for the Integrative Study of Animal Behavior Laboratory (CISAB Lab) for advice. S.K. thanks the staff of laboratory animal resources (LAR) of IU for their assistance in maintaining her mice colony.

## Author contributions

S.K conceptualized and led the study. S.K., A.P., and M.K. performed the *in vivo* mouse experiments. S.K. performed all the *in vitro* assays. H.M. conducted pS6-IP RNA-Seq experiments. A.M. analyzed the results of RNA sequencing, read the manuscript, and discussed with S.K. while finalizing it as a paper. K.D. and C.K. edited the manuscript and provided important comments on the organization and experimental approaches. H.A.S. and M.V.N. prepared SBT solution and analyzed the BCP from Sigma-Aldrich, read and edited the manuscript. S.K. wrote the manuscript with input from other authors.

## Supporting information

**S1 Fig.**
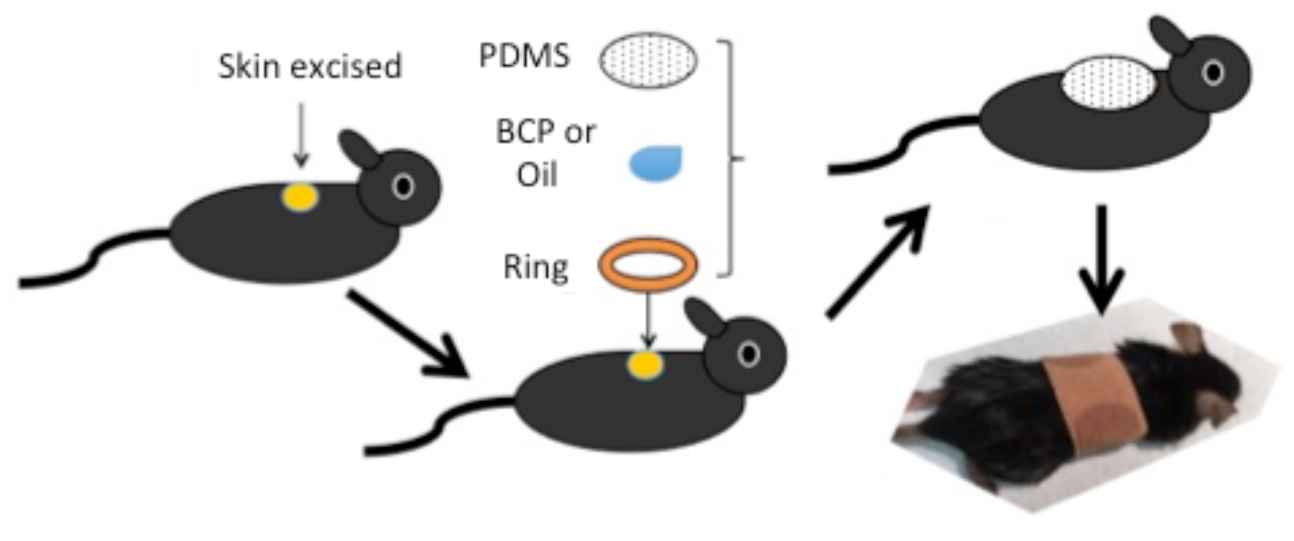
Reservoir to hold treatment media. Small (5 mm x 5 mm) full-thickness excisions were made on the backs of adult mice and methods to hold the treatment buffer solution on the wounded area were developed. A silicone ring, which held 50 μL of buffer solution, was attached to the wounded area using a liquid sealing bandage (center). After the buffer solution was placed in the ring, a transparent lid made of polydimethylsiloxane (PDMS) was installed on top of the ring and was then sealed with the liquid sealing bandage (center and upper right). A gauze bandage was used to cover the body to prevent the mouse from removing the ring (lower right).

**S2. Fig.**
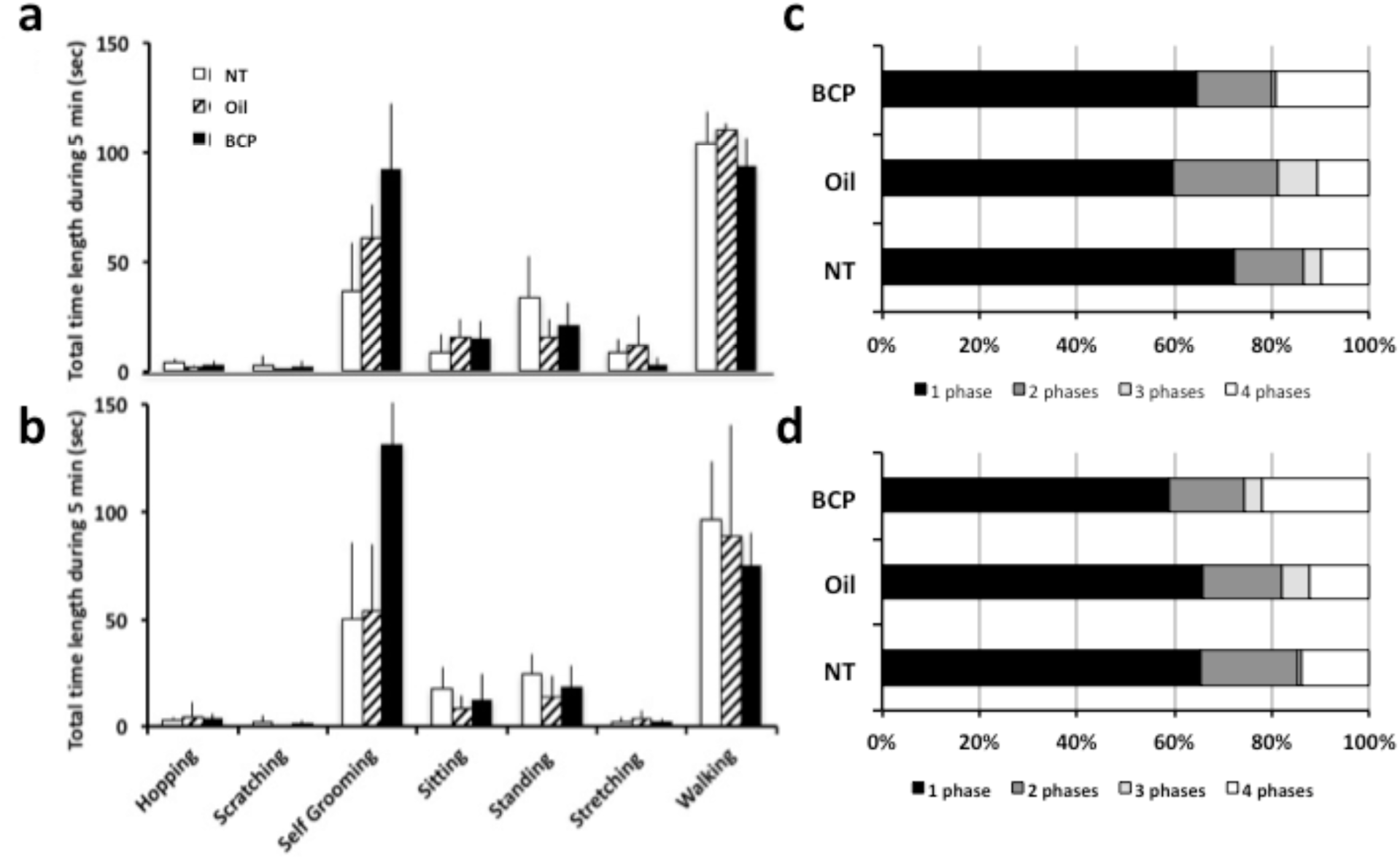
Behavioral patterns observed by BCP, Oil, and NT group. A concern in the use of BCP is that it may cause allergic responses. An oxidation product of BCP, *i.e*., caryophyllene oxide, is known as mild allergen [91]. To examine whether BCP has side effects, we conducted open-field tests on post-surgery day 1 (PS1) and 3 (PS3) and conducted behavioral analyses, specifically focusing on behavior patterns including scratching behaviors as an evidence of sensitization. In figures, NT group mice are mice without skin excision but went through all other steps and kept bandage on the same time length. There was almost no incidence of scratch behaviors on post-surgery day 1 **(a)** nor on post-surgery day 3 **(b)**. Statistically significant differences were found in self-grooming behaviors of post-surgery day 3 (ANOVA, *F*_2,15_=4.585, *P*=0.028). Self-grooming behaviors are known to increase at both high and low stress situations [92, 93]. If there are differences in the way BCP group mice did self-grooming behaviors, it could be due to BCP treatment. We classified self-grooming behaviors by the part in the body they groom and called them Phase 1 to Phase 4 following earlier studies [92, 93], and analyzed the self-grooming behaviors. We found no differences among the groups in the way self-grooming behaviors was conducted on both post-surgery day 1 and 3 (S2 Fig c,d), which suggest that BCP treatment did not cause mice to self-groom in a different way. **(c)** and **(d)** show the % of short to long, full sequences of self-grooming behaviors depending on the group on post-surgery day 1 **(c)** (NT, n=6, Oil, n=6, BCP, n=7) and 3 **(d)** (NT, n=6, Oil n=5, BCP, n=7). Classification of self-grooming behavior is as follows [92, 93]: around the nose area (Phase I), around the face (Phase II), around the head and ears (Phase III), and to the body (Phase IV). Groomings toward the bandage were excluded from Phase IV to avoid the possibility that these grooming could be intention to remove bandages. Each incidence of grooming was classified into the number of phases they include and % of short self-groomings (include only one phase) to long full self-groomings (include four phases) were calculated to determine if BCP group showed shorter self-groomings as signs of irritation stress. The % of self-grooming with four phases was higher in the BCP group but there were no statistically significant differences among groups. Based on these differences in the amount of self-grooming behaviors, we examined the travel distances and velocity of movements when they move and found that BCP group showed less travel distance and slower velocity (S3 Fig).

**S3. Fig.**
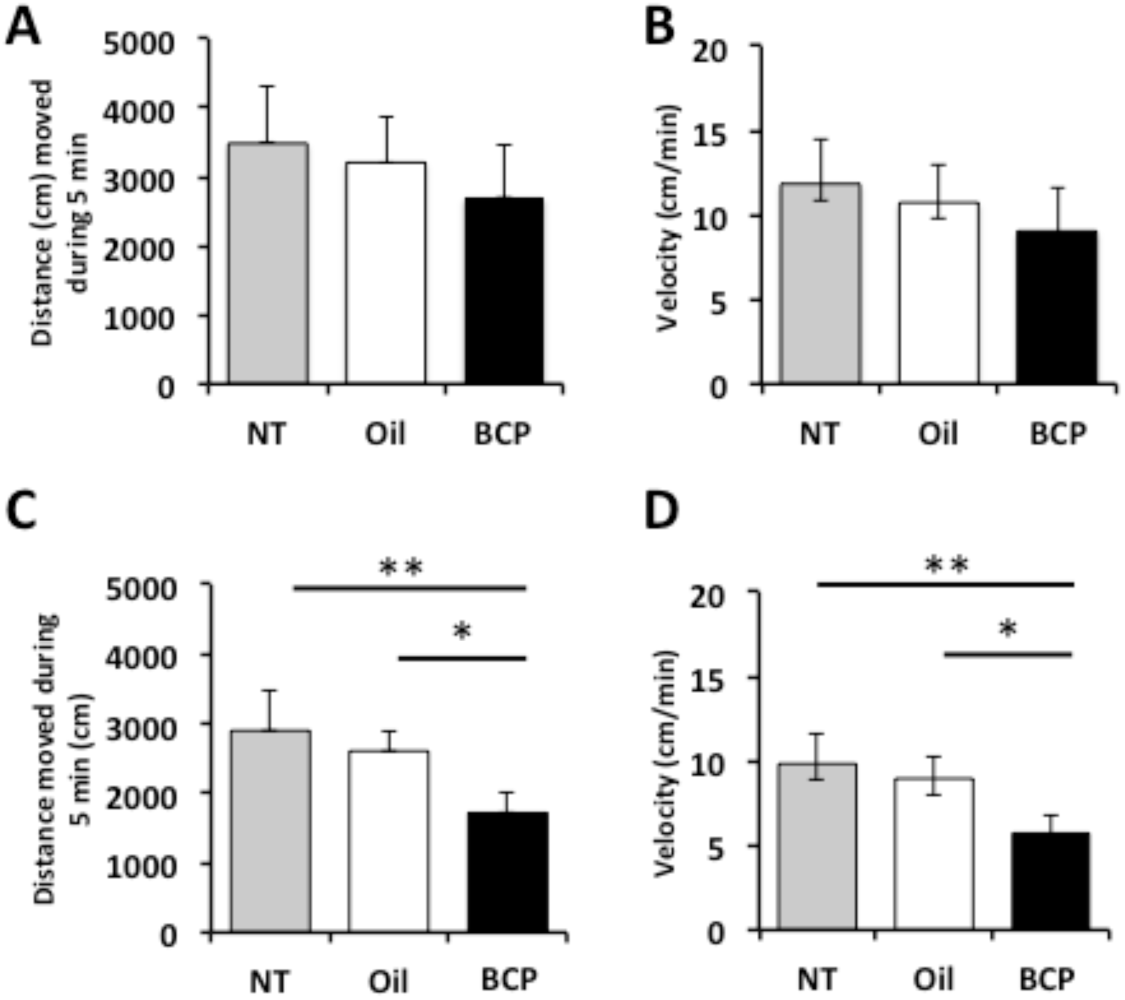
Distance traveled and velocity of movements in BCP, Oil, and NT group mice. Open-field analyses of traveling distances and moving velocity revealed that on post-surgery day 1, there were no statistically significant differences among groups in the distance traveled **(a)** and velocity of movements **(b)** (ANOVA, distance, *F*_2,21_=2.205, *P*=0.135; velocity, *F*_2,21_=2.647, *P*=0.094; NT, n=6, Oil, n=6, BCP, n=7). On post-surgery day 3, the distance traveled **(c)** was significantly shorter and the velocity was significantly slower **(d)** in the BCP group, whereas there were no differences between the Oil group and NT group (ANOVA, distance, *F*_2,9_=9.113, *P*=0.007; velocity, *F*_2,9_=9.495, *P*=0.006; NT, n=6, Oil n=5, BCP, n=7). Linalool, a chemical compound included in lavender extracts, has anxiolytic effect in mice [68]. Whether the slower movements and increased self-grooming are signs that BCP has anxiolytic influence like linalool need to be addressed in future. Overall, these results showed that the impact of BCP on behavior was the longer time staying at a place doing self-grooming and the slow movements when the mice walked, which contain no signs of irritation from allergic responses. The BCP we used contains only 1.6% of caryophyllene oxide (S4 Fig., S1 Table) and fresh BCP was applied daily. The daily change may have contributed to reduce sensitization and allergic reactions.

**S4 Fig.**
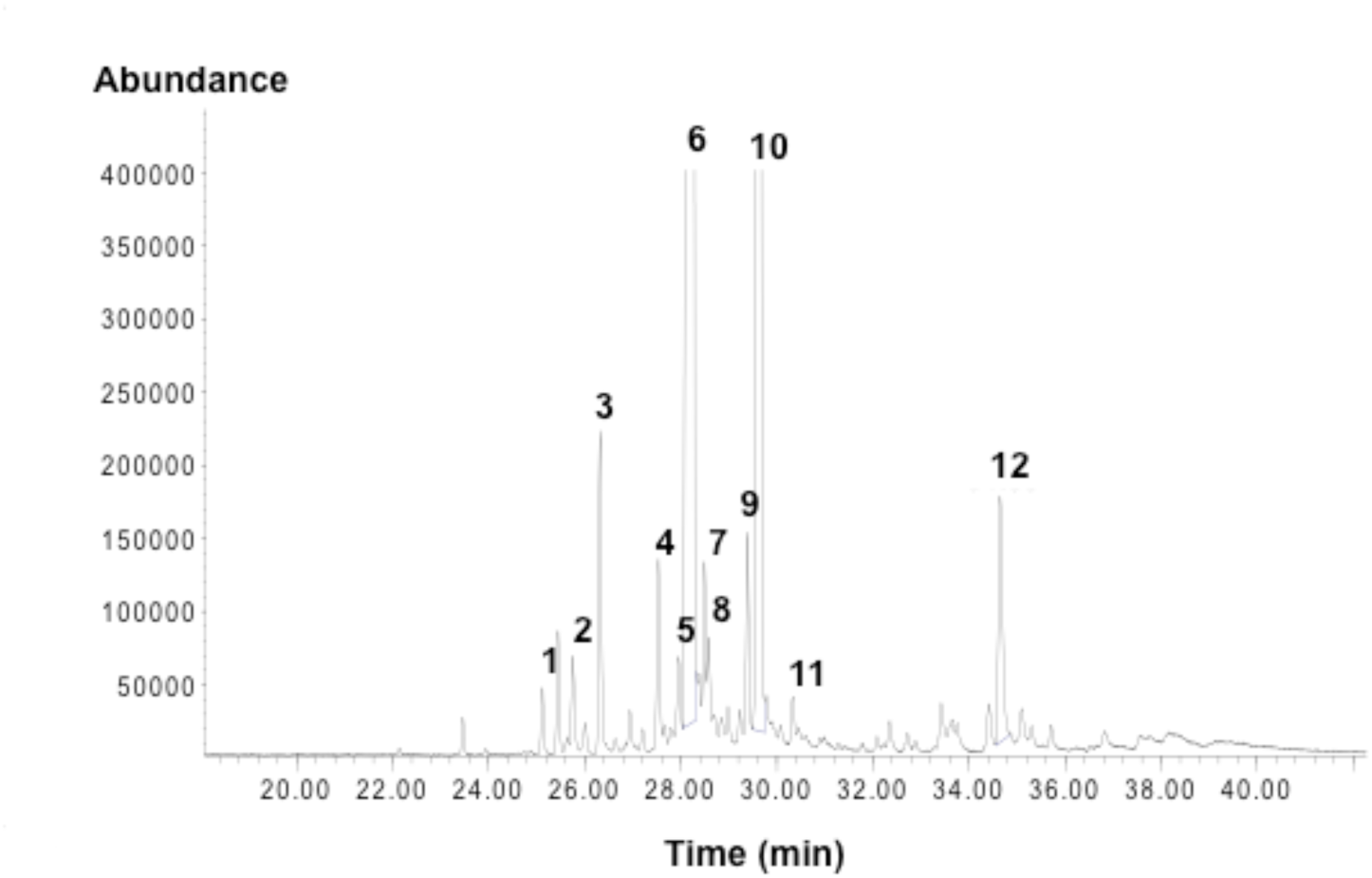
Beta-caryophyllene standard (Sigma-Aldrich) composition/GC-MS. 1: cubebene, 2, 4, 5, 7, 8: sesquiterpenes of MW 204, 3: copaene, 6: BCP, 9: neoclovene, 10: α-caryophyllene, 11: 9-epi(E)-caryophyllene, 12: caryophyllene oxide. See **S2 Table** for details.

**S5 Fig.**
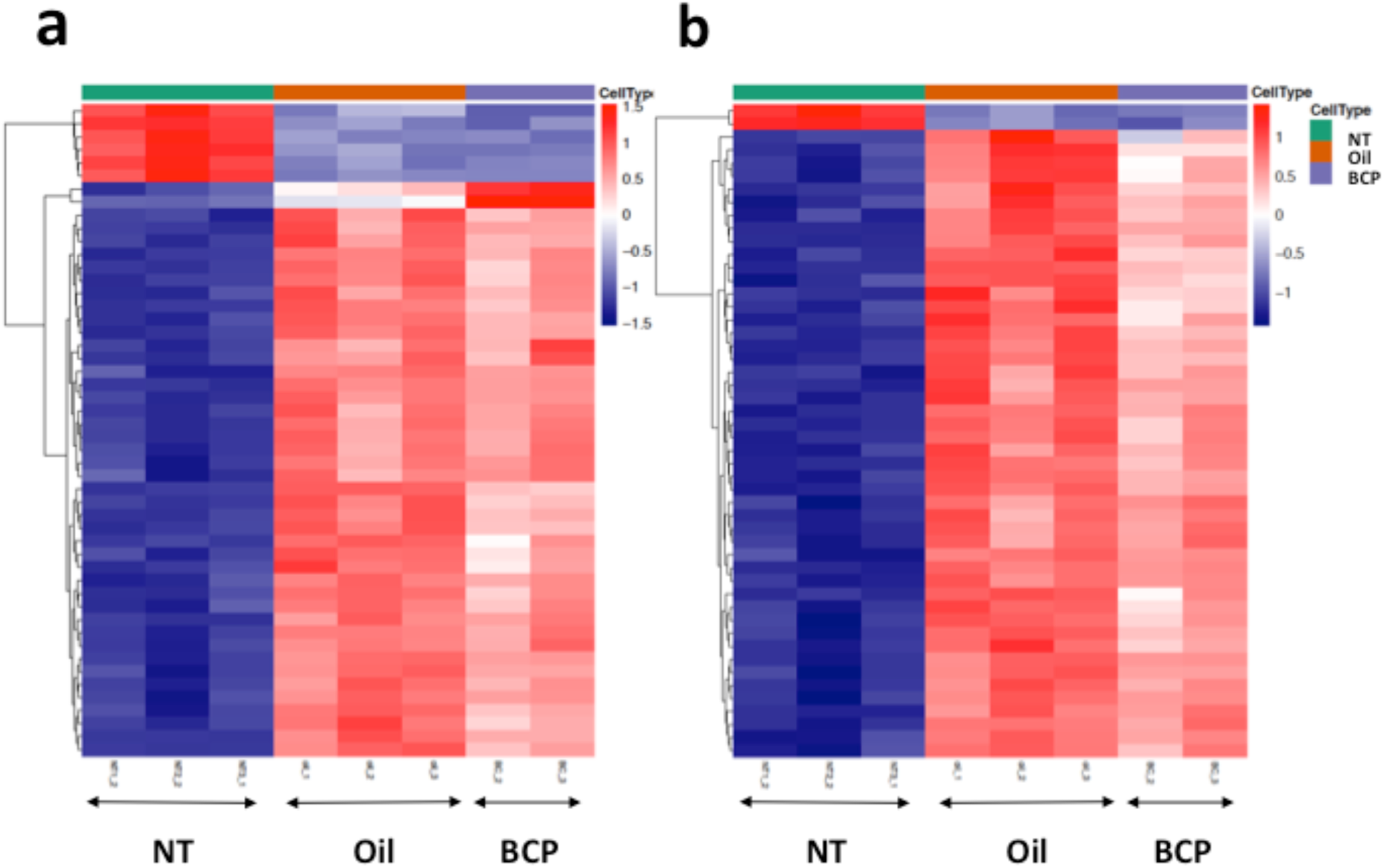
Results of RNA sequencing of post-surgery 17 hours skin and intact skin: Comparison between BCP and NT (a) and Oil and NT (b). Heatmap showing the top 50 significant gene expressions in the skin exposed to BCP (n=2) or oil (n=3), 17 to 18 hours post-surgery (inflammation stage), and in the skin of mice without skin excision (NT group) (n=3).

**S6 Fig.**
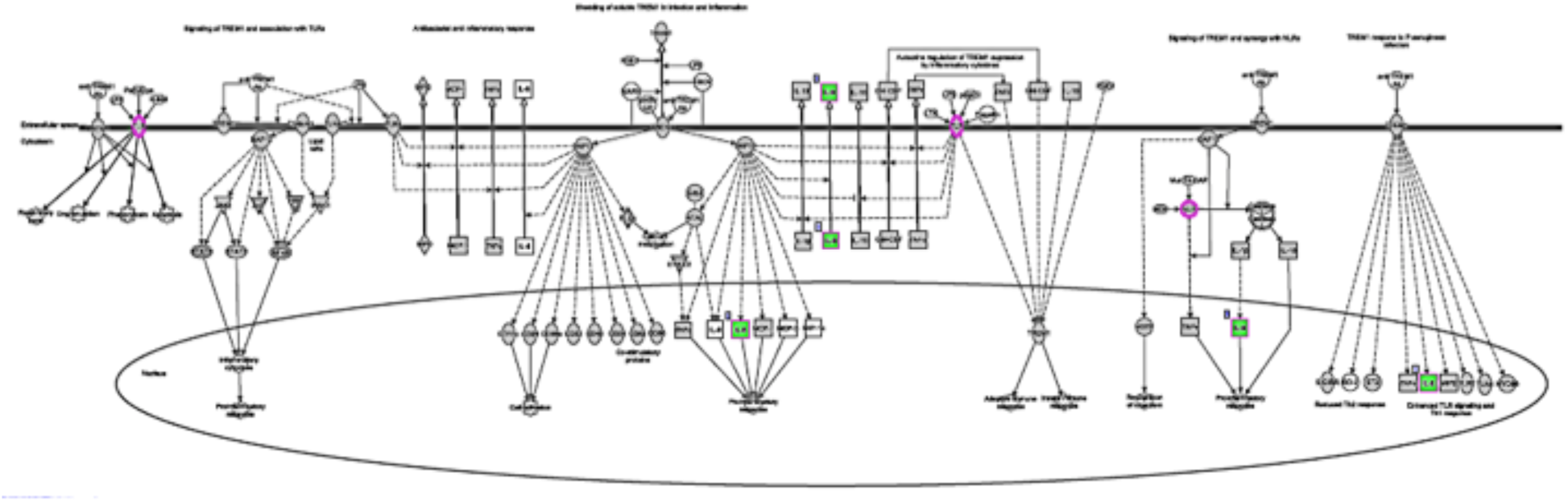
Influence of exposure to BCP on TREM1 pathway. TREM1 signaling pathway showing the genes/groups of genes up-regulated (pink) and down-regulated (green) in BCP group compared to oil group.

**S7 Fig.**
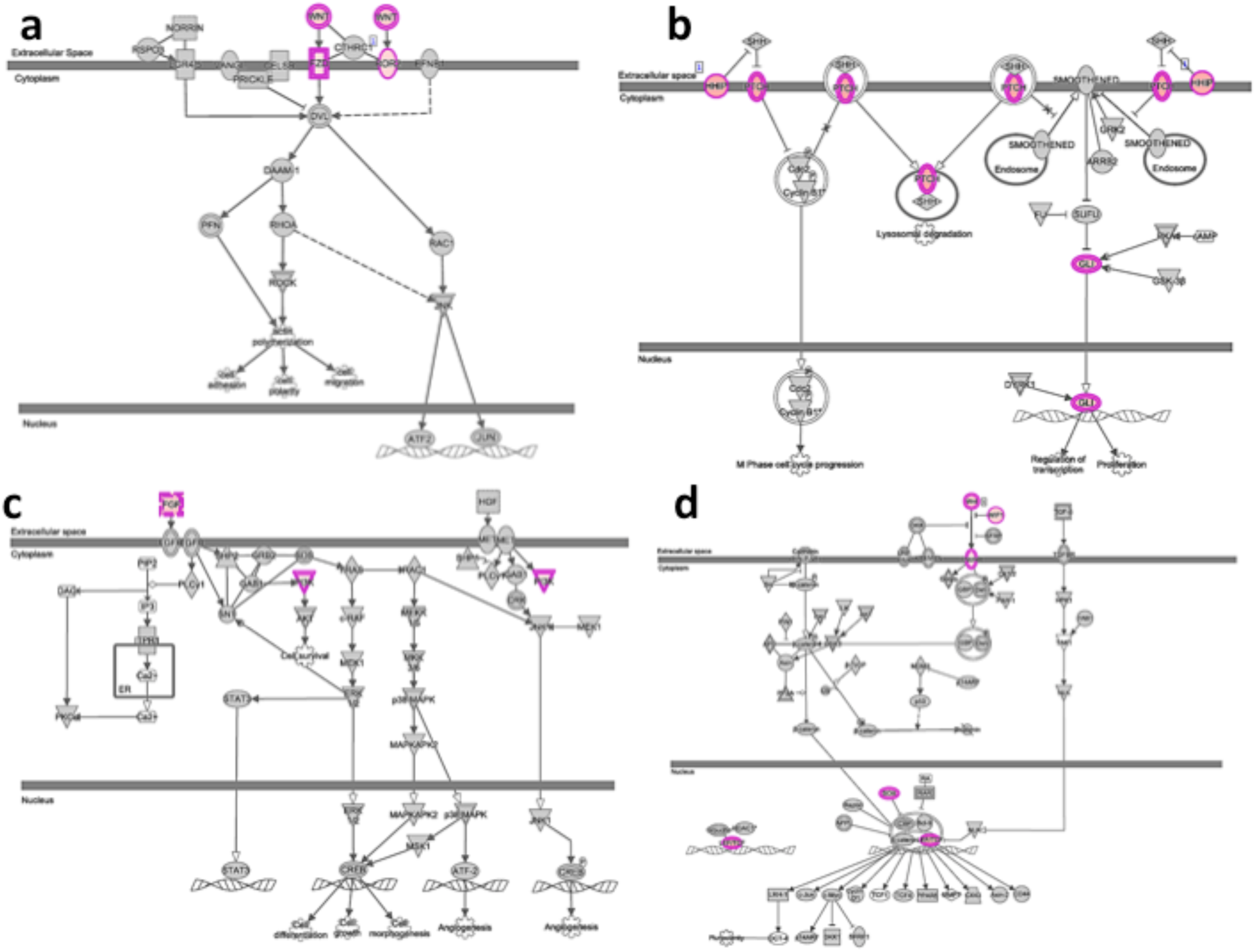
Signaling pathways showing the genes/groups of genes up-regulated (pink) in BCP group compared to oil group. (a) Sonic hedgehog signaling (shh) pathway, (b) planar cell polarity (PCP) signaling pathway, (c) fibroblast growth factor (FGF) signaling pathway, and (d) Wnt beta-catenin signaling pathway.

**S8 Fig.**
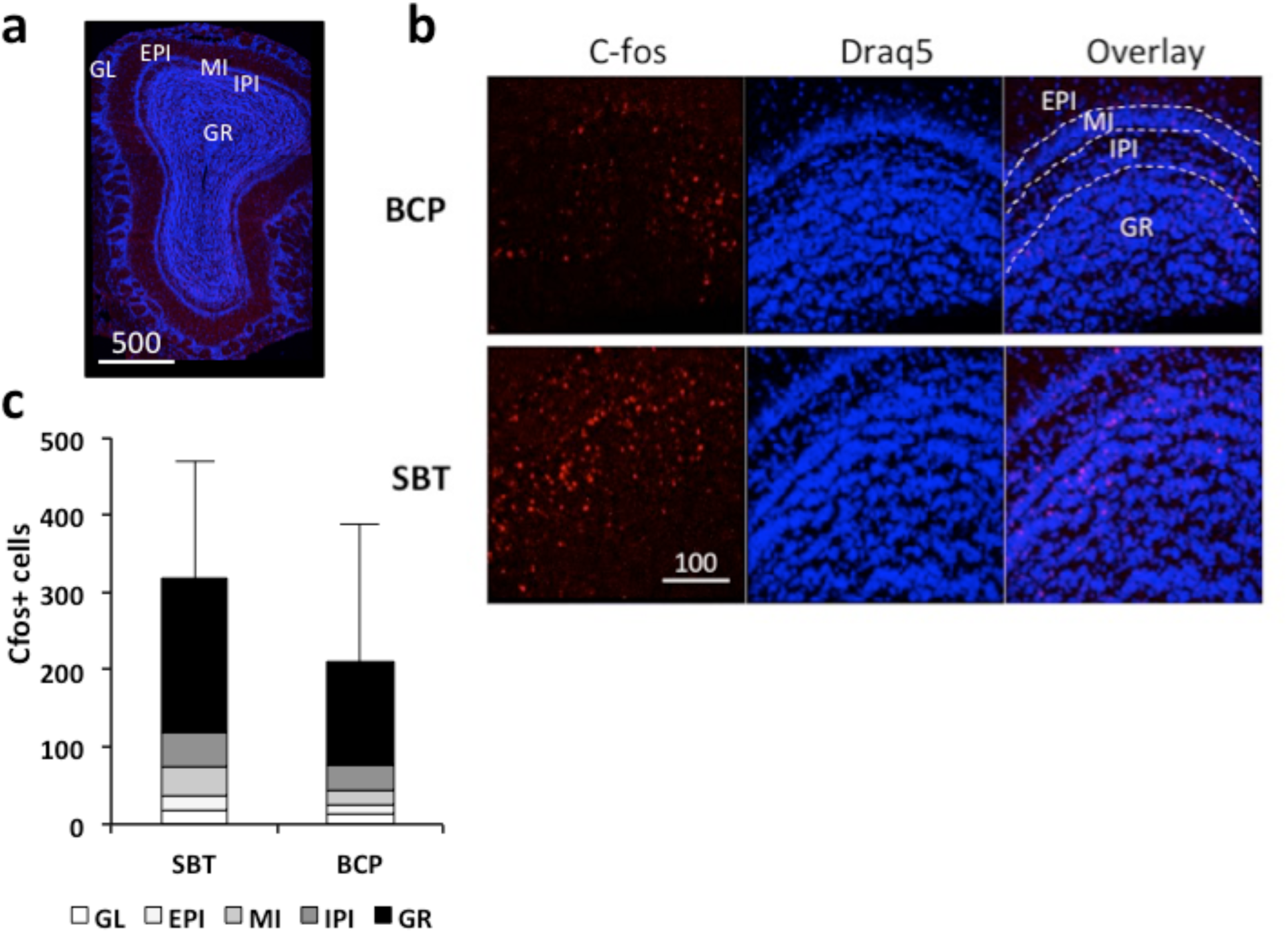
cfos expression in the olfactory bulb of mice exposed to BCP or SBT. **(a)** Location where cfos was expressed profoundly. B **(b)** To test if mice can smell BCP, intact mice were exposed to BCP (n=3) or male murine pheromone 2-sec-butyl-4,5-dihydrothiazole (SBT) (n=5) [61, 62] for 60 min and perfused. Olfactory bulb was harvested, and processed to determine c-fos protein expression. Mice exposed to BCP and SBT both showed c-fos expression in the olfactory bulb. **(c)** Numbers of cfos+ cells were not statistically significantly different between the mice exposed to SBT and BCP because of the large variance in the expression of cfos. The expression was high in the females at estrous stage. Error bars indicates sd of total number of cells. GL: glomerular layer, EPI: external plexiform layer, MI: mitral cell layer, IPI: internal plexiform layer, GR: granule cell layer.

**S9 Fig.**
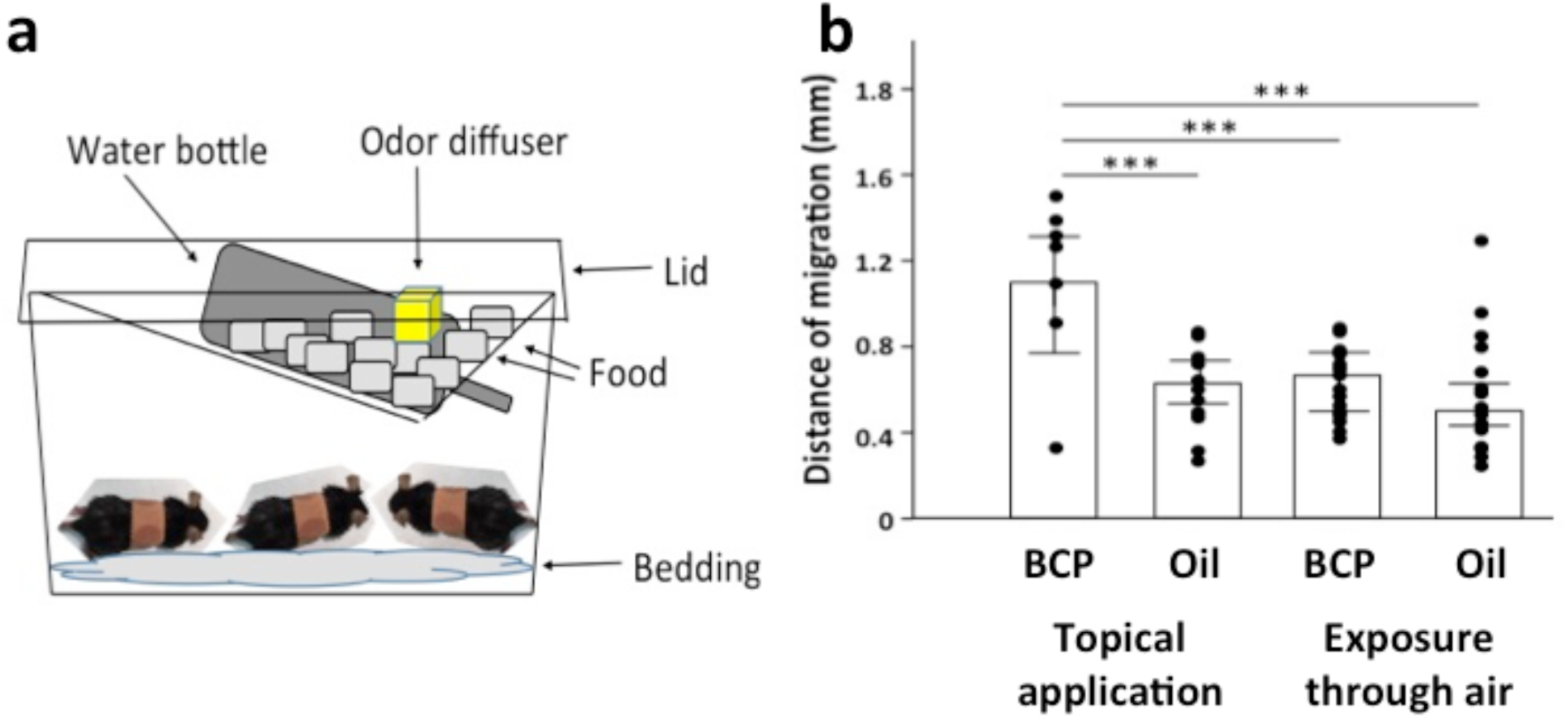
Influence of exposure to BCP through the olfactory system. Mice were exposed to BCP or olive oil (Oil) through air by putting Q-tip soaked with either BCP or olive oil in a wire mesh and placing the wire mesh on top of the food provider **(a)**. Food was placed as much as possible to avoid the possibility that mice can reach to the wire mesh. Mice went through surgery and surgery area was covered by the ring reservoir filled with olive oil and sealed by PDMS lid, which was covered by a bandage. **(b)** Exposure to BCP through air did not differ from control level and significantly different from topical application of BCP (ANOVA, BCP vs Oil F1,66=15.803, P<0.001, Topical vs Air F1,66=13.01, P=0.001, Interaction F1,66=6.701, P=0.012, BCP-Top, n=16, Oil-Top, n=16, BCP-Air, n=18, Oil-Air, n=20) ***: P<0.001, Tukey’s pairwise comparison was used for pair wise comparison.

**S1 Table.**
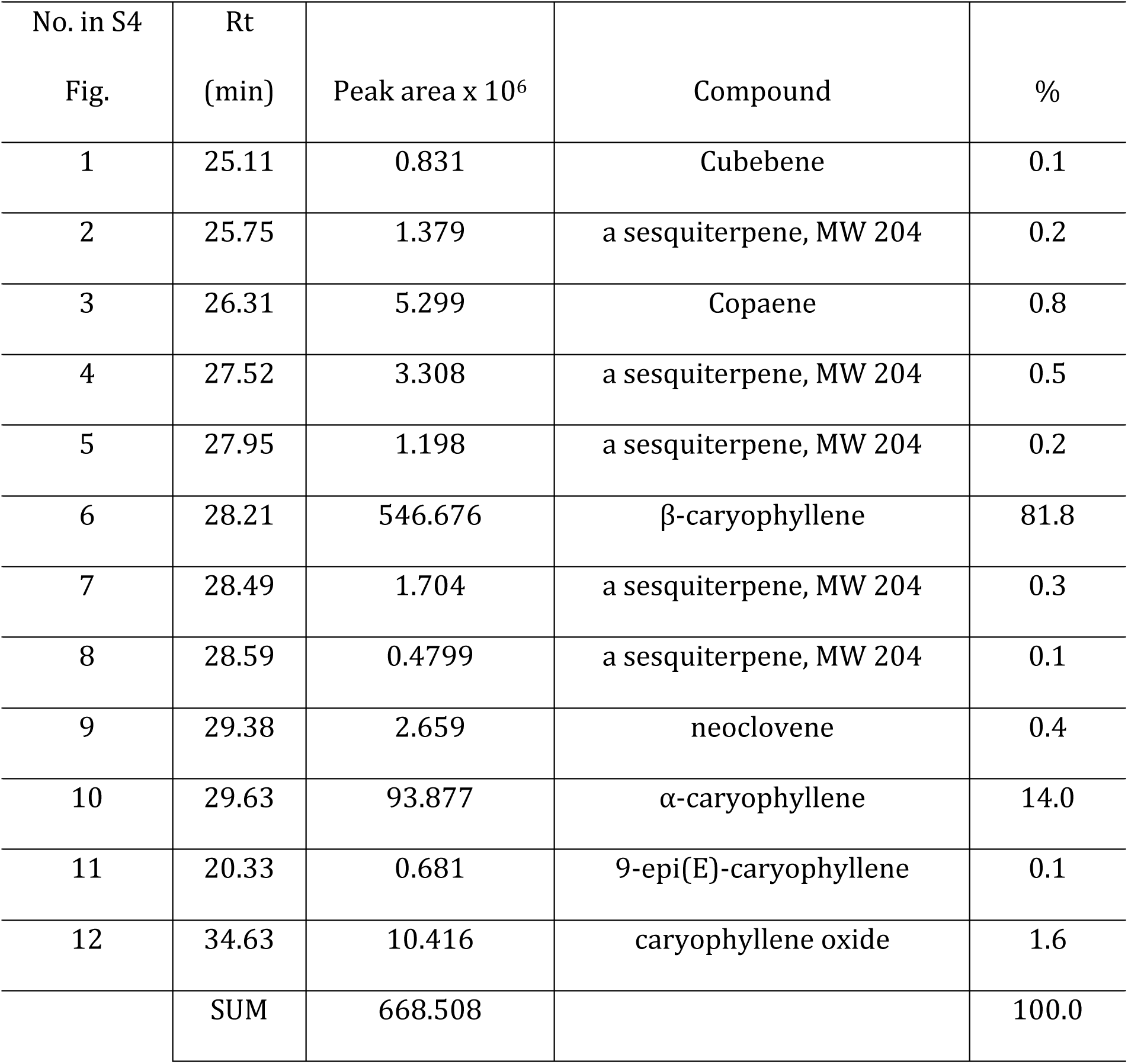
Beta-caryophyllene standard (W225207, Sigma-Aldrich) composition/GC-MS.

| No. in S4<br>Fig. | Rt<br>(min) | Peak area x 10<sup>6</sup> | Compound | % |
| --- | --- | --- | --- | --- |
| 1 | 25.11 | 0.831 | Cubebene | 0.1 |
| 2 | 25.75 | 1.379 | a sesquiterpene, MW 204 | 0.2 |
| 3 | 26.31 | 5.299 | Copaene | 0.8 |
| 4 | 27.52 | 3.308 | a sesquiterpene, MW 204 | 0.5 |
| 5 | 27.95 | 1.198 | a sesquiterpene, MW 204 | 0.2 |
| 6 | 28.21 | 546.676 | β-caryophyllene | 81.8 |
| 7 | 28.49 | 1.704 | a sesquiterpene, MW 204 | 0.3 |
| 8 | 28.59 | 0.4799 | a sesquiterpene, MW 204 | 0.1 |
| 9 | 29.38 | 2.659 | neoclovene | 0.4 |
| 10 | 29.63 | 93.877 | α-caryophyllene | 14.0 |
| 11 | 20.33 | 0.681 | 9-epi(E)-caryophyllene | 0.1 |
| 12 | 34.63 | 10.416 | caryophyllene oxide | 1.6 |
|  | SUM | 668.508 |  | 100.0 |

**S2 Table.** Olfactory receptor genes in the olfactory epithelium up-regulated after 1 hour exposure to BCP.

| Olfactory Receptor Gene | Log2FC | PValue |
| --- | --- | --- |
| *Olfr340* | 4.892 | 0.005 |
| *Olfr1480* | 1.881 | 0.028 |
| *Olfr103* | 1.656 | 0.001 |
| *Olfr1054* | 1.469 | 0.028 |
| *Olfr44* | 1.425 | 0.001 |
| *Olfr935* | 1.382 | 0.016 |
| *Olfr1152* | 1.325 | 0.041 |
| *Olfr616* | 1.246 | 0.037 |
| *Olfr1356* | 1.138 | 0.035 |
| *Olfr508* | 1.124 | 0.048 |
| *Olfr111* | 1.066 | 0.021 |
| *Olfr1258* | 1.013 | 0.002 |
| *Olfr33* | 0.928 | 0.042 |
| *Olfr874* | 0.880 | 0.026 |
| *Olfr599* | 0.810 | 0.022 |
| *Olfr92* | 0.780 | 0.046 |
| *Olfr716* | 0.736 | 0.015 |
| *Olfr701* | 0.729 | 0.048 |
| *Olfr1427* | 0.687 | 0.039 |

**S3 Table.** Olfactory receptors expressed in skin after exposure to BCP, compared with Oil group.

| Olfactory receptor genes | Log2FC | Pvalue |
| --- | --- | --- |
| *Olfr267* | 2.575 | 0.355 |
| *Olfr624* | 1.336 | 0.544 |
| *Olfr912* | 0.497 | 0.661 |
| *Olfr1396* | 0.378 | 0.796 |
| *Olfr1034* | 0.344 | 0.859 |
| *Olfr110* | 0.256 | 0.780 |
| *Olfr212* | 0.211 | 0.925 |
| *Olfr482* | 0.075 | 0.737 |
| *Olfr558* | -0.115 | 0.788 |
| *Olfr920* | -0.14 | 0.777 |
| *Olfr1392* | -0.151 | 0.965 |
| *Olfr750* | -0.222 | 0.937 |
| *Olfr733* | -0.243 | 0.929 |
| *Olfr78* | -0.249 | 0.683 |
| *Olfr523* | -0.279 | 0.862 |
| *Olfr560* | -0.296 | 0.871 |
| *Olfr77* | -0.466 | 0.668 |
| *Olfr346* | -0.81 | 0.714 |
| *Olfr56* | -0.836 | 0.264 |
| *Olfr466* | -0.847 | 0.588 |
| *Olfr1378* | -0.853 | 0.636 |
| *Olfr92* | -0.883 | 0.604 |
| *Olfr1033* | -0.898 | 0.221 |
| *Olfr111* | -1.06 | 0.134 |
| *Olfr1383* | -1.324 | 0.613 |
| *Olfr1388* | -1.376 | 0.451 |
| *Olfr433* | -1.806 | 0.405 |
| *Olfr1393* | -2.218 | 0.395 |
| *Olfr432* | -2.707 | 0.440 |
| *Olfr1564* | -2.74 | 0.265 |

## Footnotes

1 Conspecific: Technical terms in the study field of animal behavior and ethology, meaning the same species

2 Cannabinoid receptors G-protein receptors first found as the receptors for Δ^9^-tetrahydrocannabinol (THC), which is the main psychoactive constituent of *Cannabis sativa* (marijuana). Two receptors have been identified, cannabinoid receptors 1 and 2 (Zou S, Kumar, U. *Int. J. Mol. Sci*, 2018, 19, 833).

## Notes

#### Summary of Updates

Edited English. Included references recommended by the reviewers.

